# Reanalysis of deep-sequencing data from Austria points towards a small SARS-COV-2 transmission bottleneck on the order of one to three virions

**DOI:** 10.1101/2021.02.22.432096

**Authors:** Michael A. Martin, Katia Koelle

## Abstract

An early analysis of SARS-CoV-2 deep-sequencing data that combined epidemiological and genetic data to characterize the transmission dynamics of the virus in and beyond Austria concluded that the size of the virus’s transmission bottleneck was large – on the order of 1000 virions. We performed new computational analyses using these deep-sequenced samples from Austria. Our analyses included characterization of transmission bottleneck sizes across a range of variant calling thresholds and examination of patterns of shared low-frequency variants between transmission pairs in cases where *de novo* genetic variation was present in the recipient. From these analyses, among others, we found that SARS-CoV-2 transmission bottlenecks are instead likely to be very tight, on the order of 1-3 virions. These findings have important consequences for understanding how SARS-CoV-2 evolves between hosts and the processes shaping genetic variation observed at the population level.

---

In their recent research article *(1)*, Popa, Genger et al. combined epidemiological and viral genetic data to characterize the transmission dynamics of SARS-CoV-2 in Austria between February and April 2020. The genetic data they analyzed comprised >500 deep-sequenced virus samples. Beyond using consensus-level SARS-CoV-2 sequences to infer transmission clusters within Austria and to examine the role that Austria played in seeding regional epidemics elsewhere in Europe, the authors used their sequenced samples to characterize mutational dynamics within hosts and along short transmission chains. While we believe that the findings from their consensus-level genetic analysis are robust, we here revisit their analyses of mutational dynamics at the below-the-consensus level. From our reanalysis, we conclude that transmission bottleneck sizes are not on the order of 1000 virions as concluded by the authors, but instead much smaller.

Our decision to revisit Popa, Genger et al.’s conclusions on transmission bottleneck sizes stems from curious patterns present in some of their figures. First, inferred bottleneck size estimates using a 3% variant calling threshold were bimodal, with 14 of the 39 transmission pairs having an inferred bottleneck size (*N*_b_) of <10 and the remaining 25 pairs having *N*_b_ estimates of 115-5000 (their Figure S4G). Further, when a 1% variant calling threshold was used, only a single transmission pair retained an *N*_b_ estimate of <10 (their Figure 5B). In an attempt to understand these patterns, we first reanalyzed their deep sequencing data and recalled variants using their pipeline (*Supplementary Methods*). In the analyses presented below, we use these recalled variant frequencies, which appear to be highly similar to those presented in Popa, Genger et al. based on the “tv plots” published as part of their article (10.5281/zenodo.4247401).

As expected, re-estimation of transmission bottleneck sizes at variant calling thresholds of 1% and 3% yielded similar results to those shown in *(1)* (Figure S1A,B). During this analysis, we noticed that bottleneck size estimates dropped, sometimes precipitously, when going from a 1% cutoff to a 3% cutoff for every one of the 13 transmission pairs that had donors with a maximum iSNV frequency of >6% (Figure 1A; *p* = 0.004 using a paired t-test). Since increasing the variant calling threshold would remove low-frequency iSNVs from analysis, these consistent decreases in *N*_b_ estimates could come about if low-frequency donor iSNVs indicated that bottleneck sizes were large while high-frequency donor iSNVs instead indicated that bottleneck sizes were small. Examination of low-frequency iSNVs across donor-recipient pairs indeed indicate high levels of congruence between their frequencies (Figure 1B inset; Figures S2), which would suggest wide transmission bottlenecks. In contrast, high-frequency donor iSNVs rarely appeared to be transmitted to their corresponding recipient (Figures S2), suggesting narrow transmission bottlenecks.

**Figure 1.**
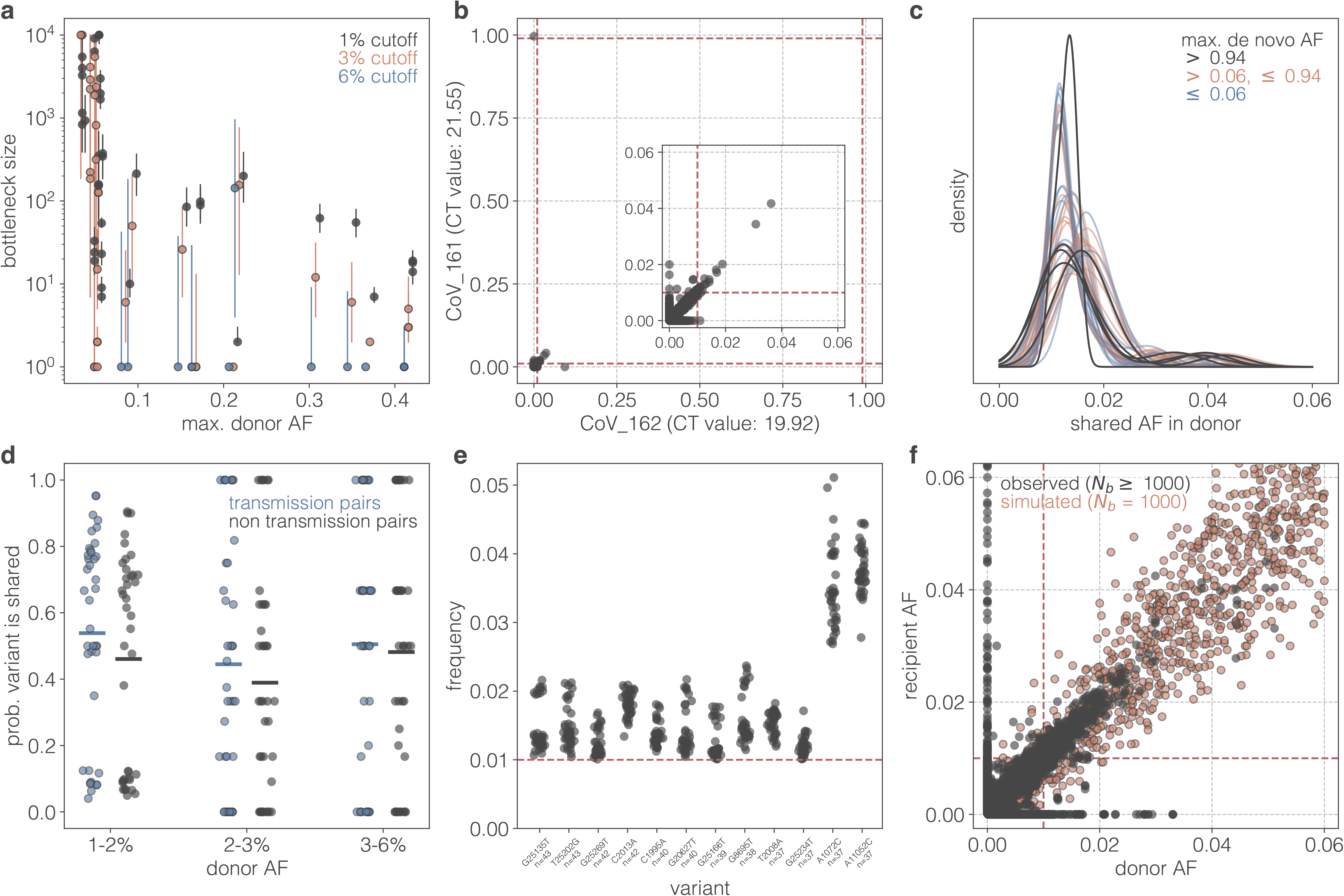
Transmission bottleneck sizes and patterns of shared viral genetic diversity between the transmission pairs studied in Popa, Genger et al. A) Bottleneck size estimates for 39 epidemiological confirmed SARS CoV-2 transmission pairs using variant calling thresholds of 1% ([0.01, 0.99]), 3% ([0.01, 0.97]), and 6% ([0.06, 0.94]). Estimates are based on all iSNVs that pass the quality filtering thresholds. Maximum likelihood estimates are indicated by a colored circle and vertical lines show 95% confidence intervals. Estimates are plotted according to the maximum iSNV frequency observed in the donor of the transmission pair. Estimates from the same transmission pair are offset slightly on the x-axis to aid in visualization. iSNV frequencies (referred to in the figures as ‘AF’ for allele frequencies) are based on variant calling relative to donor-specific reference sequences. Transmission bottleneck sizes quantify the number of virions transmitted from donor to recipient that successfully establish infection. B) All iSNVs observed in either the donor and/or the recipient of the epidemiologically confirmed CoV_162 → CoV_161 transmission pair. Donor iSNV frequency is shown on the x-axis and recipient iSNV frequency is shown on the y-axis. Note the de novo variant in the recipient (C26894U) that is fixed within individual 161 and absent from individual 162. Inset highlights low-frequency iSNVs. Red-dotted lines show the 1% variant calling threshold. iSNV frequencies are based on variant calling relative to donor-specific reference sequences. Here, all iSNVs that pass quality filtering thresholds are shown, regardless of whether they fall above or below the 1% variant calling threshold. C) Gaussian kernel density estimates of low-frequency ([0.01, 0.06]) iSNVs present in the donor that are also shared with the recipient. Lines are colored by the maximum de novo iSNV frequency in the recipient. iSNV frequencies are based on variant calling relative to donor-specific reference sequence. D) Probability that a given iSNV present in a donor is shared with a recipient host. Blue dots show probabilities for epidemiology linked pairs while black dots show probabilities for random, epidemiologically unlinked donor-recipient pairs. Random pairs are generated such that the random recipient is not a member of the same family as the focal donor or recipient and is not a known recipient of that donor sample. iSNVs are binned based on their frequency in the donor: [0.01, 0.02), [0.02, 0.03), [0.03, 0.06). Allele frequencies are based on variant calling relative to Wuhan/Hu-1. Differences between the epidemiologically-linked and –unlinked probability distributions were assessed using the Kolmogorov-Smirnov test. This test failed to find significant differences between these distributions in the 1-2% donor frequency group (p = 0.389), in the 2-3% donor frequency group (p = 0.752), and in the 3-6% frequency group (p >0.999). E) Top 12 most abundantly shared iSNVs amongst the 43 samples involved in the 39 transmission pairs. iSNVs are ordered by the number of samples in which they were found. Each dot represents the allele frequency of that iSNV in a given sample. Red-dotted line shows the 1% variant calling threshold. Allele frequencies are based on variant calling relative to Wuhan/Hu-1. F) Patterns of shared viral genetic diversity between transmission pairs with a large bottleneck. Block dots show all iSNVs observed in either the donor and/or recipient for all transmission pairs with an estimated bottleneck size of ≥1000 at a variant calling threshold of 1%. Allele frequencies are based on variant calling relative to donor-specific reference sequence. Red dots represent simulated data assuming a transmission bottleneck of 1000.

To come to terms with these conflicting patterns, we considered genetic variation that appeared *de novo* in recipient hosts. This genetic variation appears in the “tv plots” as iSNVs absent from a donor but present in a corresponding recipient. When a *de novo* variant is observed as fixed in a recipient sample, we should not observe any shared iSNVs between a donor and a recipient that are present in the recipient at subclonal (i.e., not fixed) frequencies unless within-host recombination occurred extremely rapidly or the fixed *de novo* variant arose multiple times in different genetic backgrounds. However, in the transmission pairs analyzed in Popa, Genger et al., shared subclonal iSNVs – at extremely similar frequencies - are observed in several transmission pairs where there is also a fixed de novo variant present in the recipient. The transmission pair CoV_162 → CoV_161 provides an example (Figure 1B). This means that the low-frequency iSNVs shared between CoV_162 and CoV_161 are either spurious or that they arose independently in the recipient (that is, they are homoplasies). In either case, these shared low-frequency iSNVs are highly unlikely to constitute transmitted genetic variation, and as such would need to be excluded from a transmission bottleneck analysis involving this transmission pair.

While we can only conclude that the low-frequency shared iSNVs in transmission pair CoV_162 → CoV_161 are almost certainly not shared between donor and recipient as a result of transmission, transmission pairs with *de novo* fixed variants in the recipient (here, defined as >94% in frequency), transmission pairs with *de novo* high-frequency (6-94%) variants in the recipient, and transmission pairs with only low-frequency variants (<6%) in the recipient exhibit highly similar distributions of low-frequency (1-6%) shared iSNVs (Figure 1C). The similarity between these distributions indicates that these iSNVs may be subject to the same interpretation as for CoV_162 → CoV_161. Indeed, when we calculate the probability that a low-frequency donor iSNV is observed in a corresponding recipient (at ≥1%) versus observed in an epidemiologically unlinked recipient, we find that the distribution of these probabilities are highly similar (Figure 1D). It is thus highly unlikely that these shared low-frequency iSNVs are transmitted to their corresponding recipient; if this were the case, we would expect the probability of shared variants to be higher for the corresponding recipient compared to an epidemiologically unlinked one.

Given these findings that shed doubt on low-frequency iSNVs constituting transmitted genetic variation, we decided to quantify the extent to which particular iSNVs were present across the samples used in the transmission pair analyses. We found that 5 iSNVs were present in 40 or more of the 43 samples analyzed, at frequencies that fell into a very narrow range (1%-2.2%) (Figure 1E). Many other iSNVs were also present across numerous samples (Figure 1E; Figure S3, Figure S4), with the frequencies of any particular iSNV being highly similar across the samples that it appears in. This similarity in iSNV frequencies argues against these low-frequency iSNVs being homoplasies.

Finally, a comparison between observed patterns of iSNV frequencies between donors and recipients versus those expected under large transmission bottleneck sizes as inferred in Popa, Genger et al. further argues against the transmission of the low-frequency shared iSNVs. Specifically, observed iSNV frequencies from transmission pairs with inferred bottleneck sizes of *N*_b_ ≥ 1000 show that iSNVs are present in both donor and recipient at highly similar frequencies or are observed exclusively in the donor or recipient (Figure 1F). On this figure, we overlay simulated iSNV frequencies under the assumption of a bottleneck size of *N*_b_ = 1000 (*Supplemental Methods*). Juxtaposition of the observed versus theoretically-predicted iSNV frequencies highlights an inconsistency: at *N*_b_ values of ~1000, we should expect almost all (at least 96.1%) of the iSNVs present in the donor at ≥2% to be transmitted and also observed above the variant calling threshold of 1% in the recipient. However, only 77.5% of donor iSNVs within the 2-6% frequency range are observed in the corresponding recipients at ≥1% frequency. This inconsistency indicates that the low-frequency iSNVs themselves show patterns that cannot be parsimoniously explained by large transmission bottleneck sizes.

Given these findings, we re-estimated transmission bottleneck sizes using the beta-binomial method *(2)* at a conservative variant calling threshold of 6% (Figure 1A; Figure S1C). Increasing the variant calling threshold does not bias bottleneck size estimates, but it is does increase statistical uncertainty in the estimated values. At this 6% cutoff, only 13 transmission pairs had one or more donor iSNVs remaining, such that bottleneck sizes could only be estimated for these pairs. The maximum likelihood estimate for *N*_b_ was 1 for 12 out of these 13 transmission pairs; for the remaining transmission pair (CoV_198 → CoV_230), the maximum likelihood estimate was *N*_b_ = 143 virions. This transmission pair was the only one where a donor iSNV (at a frequency of ~22%) was transmitted to a recipient but remained subclonal (at a frequency of ~17%). Since the confidence intervals around these maximum likelihood estimates were large, we also estimated an overall transmission bottleneck size using the data from these 13 transmission pairs (*Supplemental Methods*). We arrived at an estimate of a mean bottleneck size of 1.21, such that 99% of successful transmissions are expected to result from 3 or fewer virions (Figure 2).

**Figure 2.**
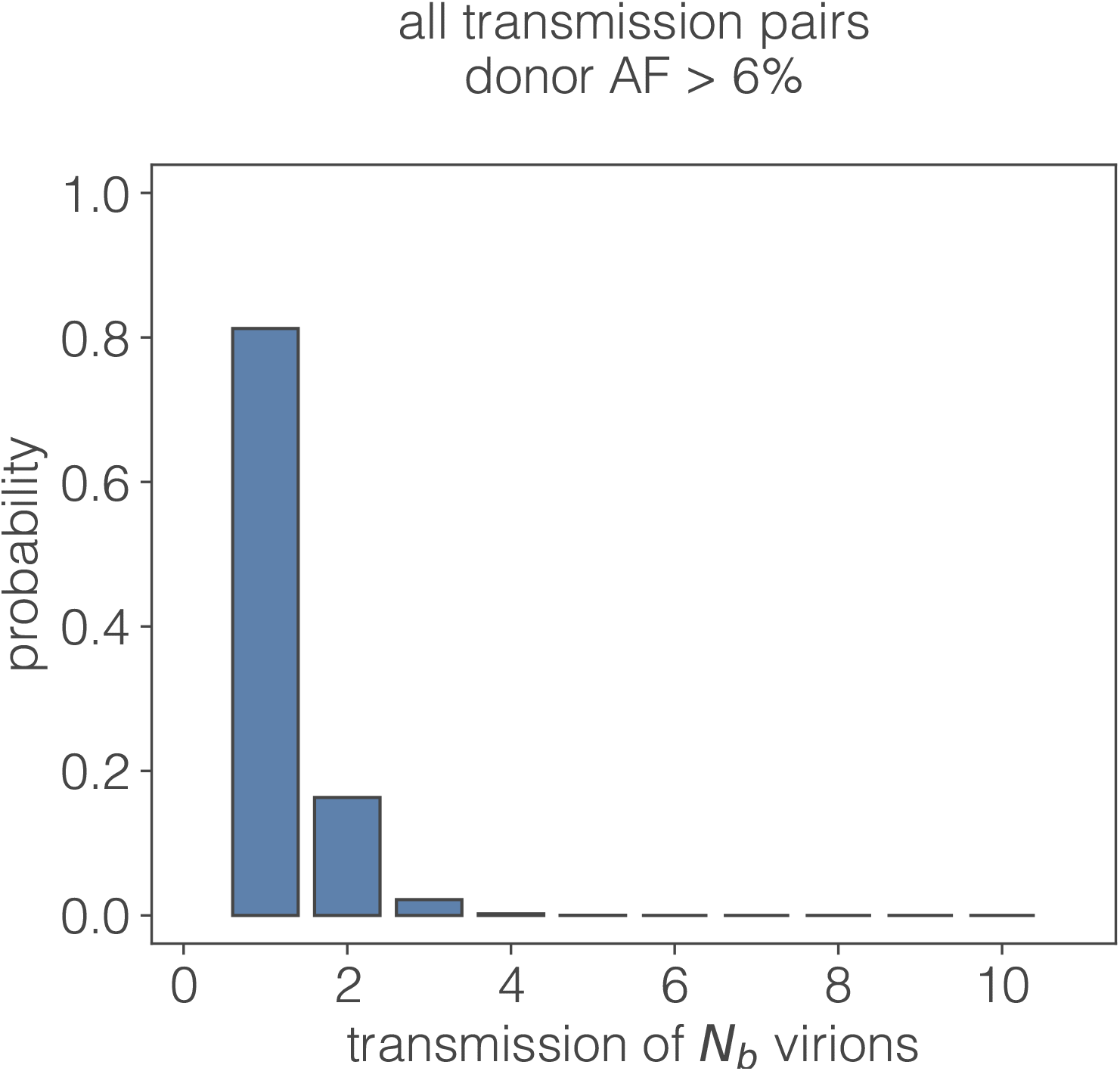
Probability of a transmission bottleneck of size *N*_*b*_ based on bottleneck size estimation using a variant-calling threshold of 6% and data from all 13 transmission pairs with one or more iSNV above this 6% threshold. The probability that a transmission involves a bottleneck size of either 1, 2, or 3 virions exceeds 99%.

Our finding of a very tight transmission bottleneck from a reanalysis of the viral deep-sequencing data from Popa, Genger et al. is consistent with conclusions from other (as yet not peer-reviewed) studies that have quantified SARS-CoV-2 transmission bottleneck sizes in humans *(3)* and other mammals *(4)*. These results indicate that SARS-CoV-2 has a narrow transmission bottleneck, similar in size to that of influenza A viruses *(5)*. Small bottleneck sizes also mean that infections generally start off with very little – if any – viral genetic diversity, such that acute infections will likely be characterized by low levels of viral diversity except in instances of superinfection, consistent with other recent (as yet not peer-reviewed) studies *(6, 7)*. Our reanalysis thus parsimoniously adds to a growing understanding of SARS-CoV-2 evolution between and within infected individuals.

## Acknowledgments

We thank Andreas Bergthaler and his group for providing clarifications on the SARS-CoV-2 deep-sequencing data submitted as part of their research article. The research reported in this technical comment was supported by National Institute of Allergy and Infectious Diseases Centers of Excellence for Influenza Research and Surveillance (CEIRS) grant HHSN272201400004C.

## References

1. A. Popa, J.-W. Genger, M. D. Nicholson, T. Penz, D. Schmid, S. W. Aberle, B. Agerer, A. Lercher, L. Endler, H. Colaço, M. Smyth, M. Schuster, M. L. Grau, F. Martínez-Jiménez, O. Pich, W. Borena, E. Pawelka, Z. Keszei, M. Senekowitsch, J. Laine, J. H. Aberle, M. Redlberger-Fritz, M. Karolyi, A. Zoufaly, S. Maritschnik, M. Borkovec, P. Hufnagl, M. Nairz, G. Weiss, M. T. Wolfinger, D. von Laer, G. Superti-Furga, N. Lopez-Bigas, E. Puchhammer-Stöckl, F. Allerberger, F. Michor, C. Bock, A. Bergthaler, Genomic epidemiology of superspreading events in Austria reveals mutational dynamics and transmission properties of SARS-CoV-2, Sci. Transl. Med. 12, eabe2555 (2020).

2. A. Sobel Leonard, D. Weissman, B. Greenbaum, E. Ghedin, K. Koelle, Transmission Bottleneck Size Estimation from Pathogen Deep-Sequencing Data, with an Application to Human Influenza A Virus, Journal of Virology, JVI.00171-17 (2017).

3. K. A. Lythgoe, M. Hall, L. Ferretti, M. de Cesare, G. MacIntyre-Cockett, A. Trebes, M. Andersson, N. Otecko, E. L. Wise, N. Moore, J. Lynch, S. Kidd, N. Cortes, M. Mori, R. Williams, G. Vernet, A. Justice, A. Green, S. M. Nicholls, M. A. Ansari, L. Abeler-Dörner, C. E. Moore, T. E. A. Peto, D. W. Eyre, R. Shaw, P. Simmonds, D. Buck, J. A. Todd, T. R. Connor, A. da Silva Filipe, J. Shepherd, E. C. Thomson, The COVID-19 Genomics UK (COG-UK) consortium, D. Bonsall, C. Fraser, T. Golubchik, Within-host genomics of SARS-CoV-2 (Genomics, 2020; http://biorxiv.org/lookup/doi/10.1101/2020.05.28.118992).

4. K. M. Braun, G. K. Moreno, P. J. Halfmann, E. B. Hodcroft, D. A. Baker, E. C. Boehm, A. M. Weiler, A. K. Haj, M. Hatta, S. Chiba, T. Maemura, Y. Kawaoka, K. Koelle, D. H. O’Connor, T. C. Friedrich, Transmission of SARS-CoV-2 in domestic cats imposes a narrow bottleneck (Microbiology, 2020; http://biorxiv.org/lookup/doi/10.1101/2020.11.16.384917).

5. J. T. McCrone, R. J. Woods, E. T. Martin, R. E. Malosh, A. S. Monto, A. S. Lauring, Stochastic processes constrain the within and between host evolution of influenza virus, eLife 7 (2018), doi:10.7554/eLife.35962.

6. G. Tonkin-Hill, I. Martincorena, R. Amato, A. R. J. Lawson, M. Gerstung, I. Johnston, D. K. Jackson, N. R. Park, S. V. Lensing, M. A. Quail, S. Gonçalves, C. Ariani, M. S. Chapman, W. L. Hamilton, L. W. Meredith, G. Hall, A. S. Jahun, Y. Chaudhry, M. Hosmillo, M. L. Pinckert, I. Georgana, A. Yakovleva, L. G. Caller, S. L. Caddy, T. Feltwell, F. A. Khokhar, C. J. Houldcroft, M. D. Curran, S. Parmar, The COVID-19 Genomics UK (COG-UK) Consortium, A. Alderton, R. Nelson, E. Harrison, J. Sillitoe, S. D. Bentley, J. C. Barrett, M. E. Torok, I. G. Goodfellow, C. Langford, D. Kwiatkowski, Wellcome Sanger Institute COVID-19 Surveillance Team, Patterns of within-host genetic diversity in SARS-CoV-2 (Genomics, 2020; http://biorxiv.org/lookup/doi/10.1101/2020.12.23.424229).

7. A. L. Valesano, K. E. Rumfelt, D. E. Dimcheff, C. N. Blair, W. J. Fitzsimmons, J. G. Petrie, E. T. Martin, A. S. Lauring, Temporal dynamics of SARS-CoV-2 mutation accumulation within and across infected hosts (Microbiology, 2021; http://biorxiv.org/lookup/doi/10.1101/2021.01.19.427330).

