## Supplementary Methods and Figures for "Reanalysis of deep-sequencing data from Austria points towards a small SARS-COV-2 transmission bottleneck on the order of one to three virions"

Aligned viral reads in the form of BAM files for all samples involved in the 39 transmission pairs described in Popa, Genger et al. *(1)* were downloaded from the National Center for Biotechnology Information (NCBI) Sequencing Read Archive (SRA) BioProject #PRJEB39849. Variants were called from these BAM files using a modified version of the variant calling pipeline used in *(1)* which was downloaded from https://github.com/Bergthalerlab/SARSCoV2_Code. Modified ARTIC (https://artic.network/) v2 primers were trimmed using iVar v. 1.3 *(2)* with a four base sliding window, no minimum quality threshold, and a minimum length of 20. Overlapping read pairs were clipped using the clipOverlap utility available in bamUtil v. 1.0.14 *(3)*. LoFreq v. 2.1.5 *(4)* was used to realign reads to correct mapping errors with the viterbi function and to insert indel quality scores using the indelqual function. Finally, LoFreq was used to call variants relative to Wuhan/Hu-1 (NC_045512.2) *(5)* requiring a minimum of 75 reads. Variant positions with a phred quality score < 90 were marked with LoFreq filter and indels with with an HRUN > 4 were marked with BCFtools v. 1.10.2 *(6)* (using HTSlib v. 1.10.2) filter. BCFtools norm was used to left-align and normalize indels and split bi-allelic records. The variant calling pipeline was run in Python v. 3.7.3 *(7)* using PyPiper v. 0.12.1 *(8)*.

For each donor sample, BCFtools view and BCFtools consensus were used to generate a donor-specific reference sequence based on the Wuhan/Hu-1 reference and any variants identified with an allele frequency > 50%. SAMtools v. 1.10 *(6)* was used to sort the viral BAM files by name and generate FASTQ files. FASTQ files for each transmission pair were realigned to the donor specific reference sequence using bwa mem v. 0.7.17-r1188 *(9)* with a seed length of 17, looking for internal seeds longer than 17*1.25, and marking shorter split hits as secondary as in *(1)*. The same pipeline as above was used to recall variants relative to the donor reference sequences.

For each transmission pair we identified the variants present in the donor at frequencies of ≥1%, ≥3% and ≥6%. These donor allele frequencies and the corresponding recipient allele frequencies were used to estimate the transmission bottleneck as described in *(10)* using MATLAB R2020A. Estimated transmission bottleneck sizes at these variant calling thresholds are shown in Figure S1.

For each transmission pair, we calculated the probability density function of low frequency ([0.01, 0.06]) variants using the guassian_kde function of Scipy v. 1.5.4 *(11)* on the donor allele frequency of all variants shared between donor and recipient. Transmission pairs are categorized based on the maximum *de novo* (i.e. present in less than 0.0001 of the reads in the donor) allele frequency in the recipient. Probability density functions of shared low-frequency variants are shown in Figure 1C.

We further calculated the transmission for these low frequency alleles by first identifying all variants present in donor samples at the following frequencies: [0.01, 0.02), [0.02, 0.03), and [0.03, 0.06) and then identifying the proportion of these variants that were identified in the recipient samples (at a frequency ≥0.01). To generate a comparison distribution, we sampled non-epidemiologically linked (i.e. not a known recipient of the focal donor sample and not in the same family as the focal donor or recipient) "recipients" for each donor sample without replacement and calculated the proportion of shared variants belonging to each category. The proportions of shared variants between transmission pairs, as well as between epidemiological unlinked samples, are shown in Figure 1D.

To compare patterns of observed iSNV frequencies against those expected under a bottleneck size of 1000 (Figure 1F), we first generated a large set (n = 1000) of low-frequency donor iSNVs by drawing their frequencies from a uniform distribution of 0 to 6%. For each of these donor iSNVs, we create a stochastic realization of the iSNV's frequency in the recipient based on the forward model on which the beta-binomial method that estimates transmission bottleneck sizes is based. Specifically, for each iSNV, we first drew a random variable $k$ from a binomial distribution with $N_b$ = 1000 number of trials and success probability given by the donor iSNV frequency. This random variable $k$ gives a stochastic realization of the number of virions in the founding population size of $N_b$ that carrying the variant allele. We then determined the frequency of the iSNV in the recipient by drawing a random variable from a beta distribution with parameters $k$ and ($N_b$ – $k$). Following this procedure for each of the 1000 iSNVs gives rise to the data points shown in orange in Figure 1F.

Overall transmission bottleneck sizes were estimated based on the assumption that transmission bottleneck sizes were distributed according to a zero-truncated Poisson-distribution. To arrive at a maximum likelihood estimate for the mean Nb, given the available iSNV frequency data for the 13 transmission pairs that had donor iSNVs that exceeded 6%, we first use the beta-binomial method to infer, for each of the 13 transmission pairs, log-likelihood values for each bottleneck size from 1 to 50. We then considered a broad range of $\lambda$ values (0 to 10) for this Poisson distribution. At any value of $\lambda$, the mean $N_b$ is given by $\frac{\lambda}{(1-e^{-\lambda})}$ . At each value of $\lambda$ considered, we calculated the probability mass of $k$ virions initiating an infection, with $p_k = \frac{\lambda^k}{(e^{\lambda}-1)k!}$ , and $k$ = 1, 2, 3, etc. The likelihood of any given value of $\lambda$ was evaluated via the following equation:

$$L(\lambda) = \sum_{n=1}^{13} \sum_{k=1}^{\infty} p_k e^{logL_{n,k}}$$

where $n$ indexes the transmission pairs and $k$ indexes the number of virions initiating infection. $logL_{n,k}$ is the log-likelihood of a bottleneck size of $k$ in transmission pair $n$. The likelihood of $\lambda$ peaked at a value of 0.042, corresponding to a mean $N_b$ of 1.21. Figure 2 shows the probability mass function of the zero-truncated Poisson distribution parameterized with a $\lambda$ of 0.042.

Two out of the 13 transmission pairs have donor samples with an unusually large number of high-frequency iSNVs. These are donor CoV_187 and donor Cov_273 (see Figure S2). Both of these donors also have high CT values (>34). Due to the possibility of transmission pairs associated with these donors biasing the mean Nb to a low value, we re-estimated the mean $N_b$ again after removing these two transmission pairs. Based on the remaining 11 transmission pairs, we find that the maximum likelihood estimate for $\lambda$ = 0.659, for the mean $N_b$ = 1.37, yielding a similar conclusion that over 99% of successful SARS-CoV-2 infections result from 3 or fewer virions.

Unless otherwise noted data was analyzed in Python v. 3.9.1 *(12)* using Pandas v. 1.1.4 *(13)* and Numpy v. 1.19.4 *(14)* and all figures were generated using MatPlotLib v. 3.3.3 *(15)*. All code to replicate the analysis is available at https://github.com/m-a-martin/sarscov2_nb_reanalysis. All data are also available at this repository, accessible in the supplementary data from *(1)*, or accessible from NCBI SRA BioProject # PRJEB39849.

**Supplemental References**


1. A. Popa, J.-W. Genger, M. D. Nicholson, T. Penz, D. Schmid, S. W. Aberle, B. Agerer, A. Lercher, L. Endler, H. Colaço, M. Smyth, M. Schuster, M. L. Grau, F. Martínez-Jiménez, O. Pich, W. Borena, E. Pawelka, Z. Keszei, M. Senekowitsch, J. Laine, J. H. Aberle, M. Redlberger-Fritz, M. Karolyi, A. Zoufaly, S. Maritschnik, M. Borkovec, P. Hufnagl, M. Nairz, G. Weiss, M. T. Wolfinger, D. von Laer, G. Superti-Furga, N. Lopez-Bigas, E. Puchhammer-Stöckl, F. Allerberger, F. Michor, C. Bock, A. Bergthaler, Genomic epidemiology of superspreading events in Austria reveals mutational dynamics and transmission properties of SARS-CoV-2, *Sci. Transl. Med.* **12**, eabe2555 (2020).

2. N. D. Grubaugh, K. Gangavarapu, J. Quick, N. L. Matteson, J. G. De Jesus, B. J. Main, A. L. Tan, L. M. Paul, D. E. Brackney, S. Grewal, N. Gurfield, K. K. A. Van Rompay, S. Isern, S. F. Michael, L. L. Coffey, N. J. Loman, K. G. Andersen, An amplicon-based sequencing framework for accurately measuring intrahost virus diversity using PrimalSeq and iVar, *Genome Biol* **20**, 8 (2019).

3. G. Jun, M. K. Wing, G. R. Abecasis, H. M. Kang, An efficient and scalable analysis framework for variant extraction and refinement from population-scale DNA sequence data, *Genome Res.* **25**, 918–925 (2015).

4. A. Wilm, P. P. K. Aw, D. Bertrand, G. H. T. Yeo, S. H. Ong, C. H. Wong, C. C. Khor, R. Petric, M. L. Hibberd, N. Nagarajan, LoFreq: A sequence-quality aware, ultra-sensitive variant caller for uncovering cell-population heterogeneity from high-throughput sequencing datasets, *Nucleic Acids Research* **40**, 11189–11201 (2012).

5. F. Wu, S. Zhao, Y.-M. Chen, W. Wang, Z.-G. Song, Y. Hu, Z.-W. Tao, J.-H. Tian, Y.-Y. Pei, M.-L. Yuan, Y.-L. Zhang, F.-H. Dai, Y. Liu, Q.-M. Wang, J.-J. Zheng, L. Xu, E. C. Holmes, Y.-Z. Zhang, A new coronavirus associated with human respiratory disease in China, *Nature* **579**, 265–269 (2020).

6. H. Li, A statistical framework for SNP calling, mutation discovery, association mapping and population genetical parameter estimation from sequencing data, *Bioinformatics* **27**, 2987–2993 (2011).

7. *Python Language Reference, version 3.7* (Python Software Foundation; http://www.python.org).

8. *PyPiper* (Databio; http://pypiper.databio.org/en/latest/faq/).

9. L. Heng, *BWA-MEM* (; http://bio-bwa.sourceforge.net).

10. A. Sobel Leonard, D. B. Weissman, B. Greenbaum, E. Ghedin, K. Koelle, D. S. Lyles, Ed. Transmission Bottleneck Size Estimation from Pathogen Deep-Sequencing Data, with an Application to Human Influenza A Virus, *Journal of Virology* **91** (2017), doi:10.1128/JVI.00171-17.

11. SciPy 1.0 Contributors, P. Virtanen, R. Gommers, T. E. Oliphant, M. Haberland, T. Reddy, D. Cournapeau, E. Burovski, P. Peterson, W. Weckesser, J. Bright, S. J. van der Walt, M. Brett, J. Wilson, K. J. Millman, N. Mayorov, A. R. J. Nelson, E. Jones, R. Kern, E. Larson, C. J. Carey, İ. Polat, Y. Feng, E. W. Moore, J. VanderPlas, D. Laxalde, J. Perktold, R. Cimrman, I. Henriksen, E. A. Quintero, C. R. Harris, A. M. Archibald, A. H. Ribeiro, F. Pedregosa, P. van Mulbregt, SciPy 1.0: fundamental algorithms for scientific computing in Python, *Nat Methods* **17**, 261–272 (2020).

12. *Python Language Reference, version 3.9* (Python Software Foundation; http://www.python.org).

13. *Pandas* (The pandas development team; https://pandas.pydata.org).

14. C. R. Harris, K. J. Millman, S. J. van der Walt, R. Gommers, P. Virtanen, D. Cournapeau, E. Wieser, J. Taylor, S. Berg, N. J. Smith, R. Kern, M. Picus, S. Hoyer, M. H. van Kerkwijk, M. Brett, A. Haldane, J. F. del Río, M. Wiebe, P. Peterson, P. Gérard-Marchant, K. Sheppard, T. Reddy, W. Weckesser, H. Abbasi, C. Gohlke, T. E. Oliphant, Array programming with NumPy, *Nature* **585**, 357–362 (2020).

15. J. D. Hunter, Matplotlib: A 2D graphics environment, *Computing in Science and Engineering* **9**, 99–104 (2007).

### Supplemental Figure Legends

**Figure S1.** Transmission bottleneck size estimates for each of the 39 transmission pairs analyzed in Popa, Genger, et al. Bottleneck sizes were estimated using A) a 1% variant calling threshold [0.01 0.99], B) a 3% variant calling threshold [0.03 0.97], and C) a 6% variant calling threshold [0.06 0.94]. Frequencies of iSNVs are based on variant calling relative to donor-specific reference sequences. Color coding of transmission pairs is as in Popa, Genger, et al. for ease of comparison. Maximum likelihood estimates are indicated by a colored circle and vertical lines show the 95% confidence intervals. (We allowed a maximum bottleneck size of 10,000 virions, while Genger, Popa, et al. allowed a maximum bottleneck size of 5,000 virions, but this difference was inconsequential to the interpretation of the results.) Note that in (B), we do not have transmission bottleneck size estimates for pairs with individual 146 as donor because the maximum iSNV frequency we found in this sample was 0.0299.

**Figure S2.** All iSNVs observed in either donor and/or recipient of all 39 epidemiologically confirmed transmission pairs. Donor iSNV frequency is shown on the x-axis and recipient iSNV frequency is shown on the y-axis. Low-frequency iSNVs are highlighted in the panel on the right.

Red-dashed lines show the 1% variant calling threshold. iSNV frequencies are based on variant calling relative to donor-specific reference sequences. CT values associated with the donor and recipient samples are provided when available. Note that high CT samples tend to have a large number of high-frequency iSNVs, while low CT samples rarely have iSNV frequencies that exceed 6%.

**Figure S3.** Allele frequencies for all iSNVs observed in at least 3 of the 43 samples involved in the 39 transmission pairs. iSNVs are ordered by the number of samples they are found in. Each dot shows the allele frequency of that iSNV in a given sample. Red-dotted lines show the 1% variant calling threshold. Allele frequencies are based on variant calling relative to Wuhan/Hu-1.

**Figure S4.** Abundance (number of samples) of all iSNVs observed in at least 3 of the 43 samples involved in the 39 transmission pairs. Each dots shows a unique iSNV, which are ordered by decreasing abundance.

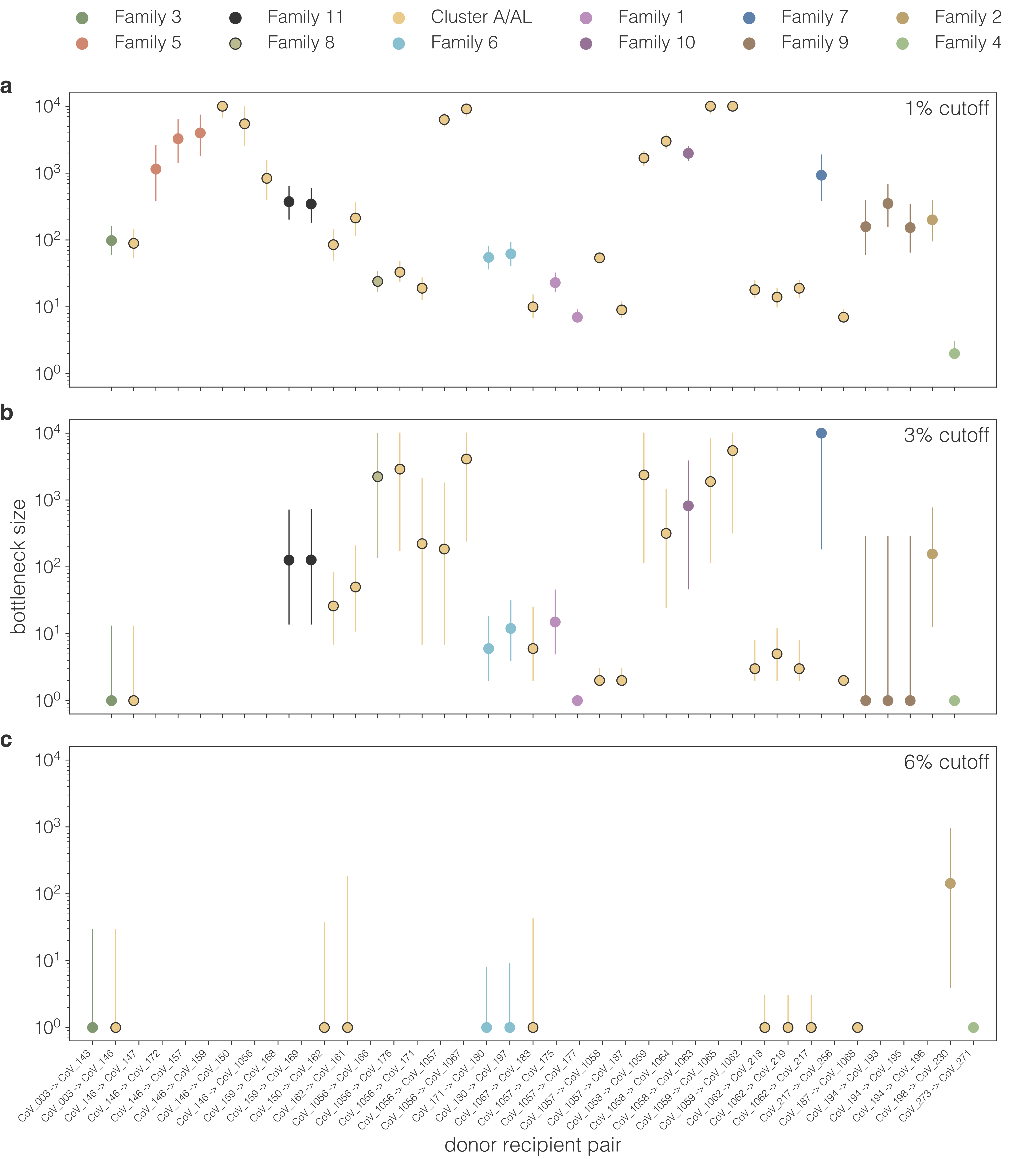

Family 3
Family 5
Family 11
Family 8
Cluster A/AL
Family 6
Family 1
Family 10
Family 7
Family 9
Family 2
Family 4
a
1% cutoff
b
3% cutoff
c
6% cutoff
bottleneck size
$10^0$
$10^1$
$10^2$
$10^3$
$10^4$
CoV_003 -> CoV_143
CoV_003 -> CoV_146
CoV_146 -> CoV_147
CoV_146 -> CoV_172
CoV_146 -> CoV_157
CoV_146 -> CoV_159
CoV_146 -> CoV_150
CoV_146 -> CoV_1056
CoV_159 -> CoV_168
CoV_159 -> CoV_169
CoV_150 -> CoV_162
CoV_162 -> CoV_161
CoV_1056 -> CoV_166
CoV_1056 -> CoV_176
CoV_1056 -> CoV_171
CoV_1056 -> CoV_1057
CoV_1056 -> CoV_1067
CoV_171 -> CoV_180
CoV_180 -> CoV_197
CoV_1067 -> CoV_183
CoV_1057 -> CoV_175
CoV_1057 -> CoV_177
CoV_1057 -> CoV_1058
CoV_1057 -> CoV_187
CoV_1058 -> CoV_1059
CoV_1058 -> CoV_1064
CoV_1058 -> CoV_1063
CoV_1059 -> CoV_1065
CoV_1059 -> CoV_1062
CoV_1062 -> CoV_218
CoV_1062 -> CoV_219
CoV_1062 -> CoV_217
CoV_217 -> CoV_256
CoV_187 -> CoV_1068
CoV_194 -> CoV_193
CoV_194 -> CoV_195
CoV_194 -> CoV_196
CoV_198 -> CoV_230
CoV_273 -> CoV_271
donor recipient pair

**aa**

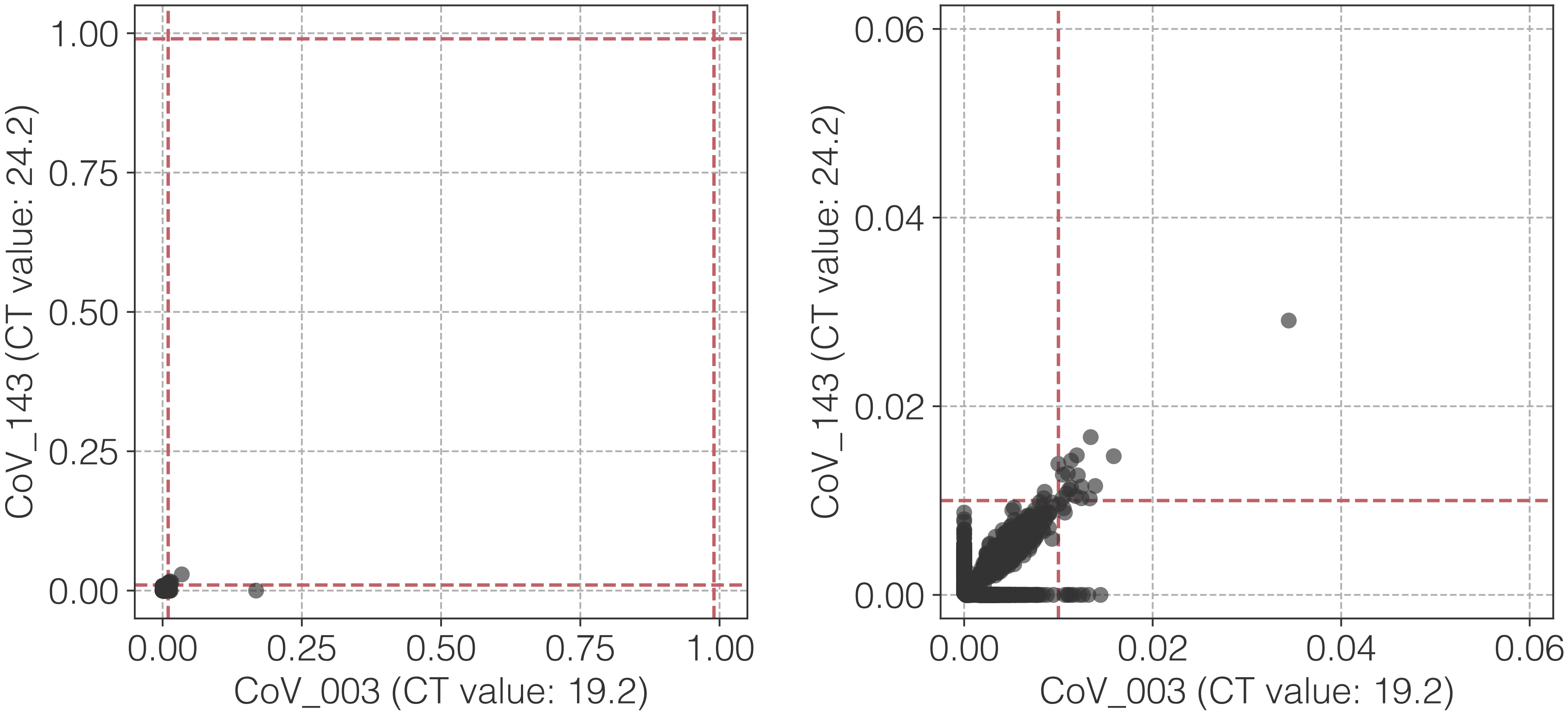

1.00
0.75
0.50
0.25
0.00
CoV_143 (CT value: 24.2)
0.00
0.25
0.50
0.75
1.00
CoV_003 (CT value: 19.2)
0.06
0.04
0.02
0.00
CoV_143 (CT value: 24.2)
0.00
0.02
0.04
0.06
CoV_003 (CT value: 19.2)

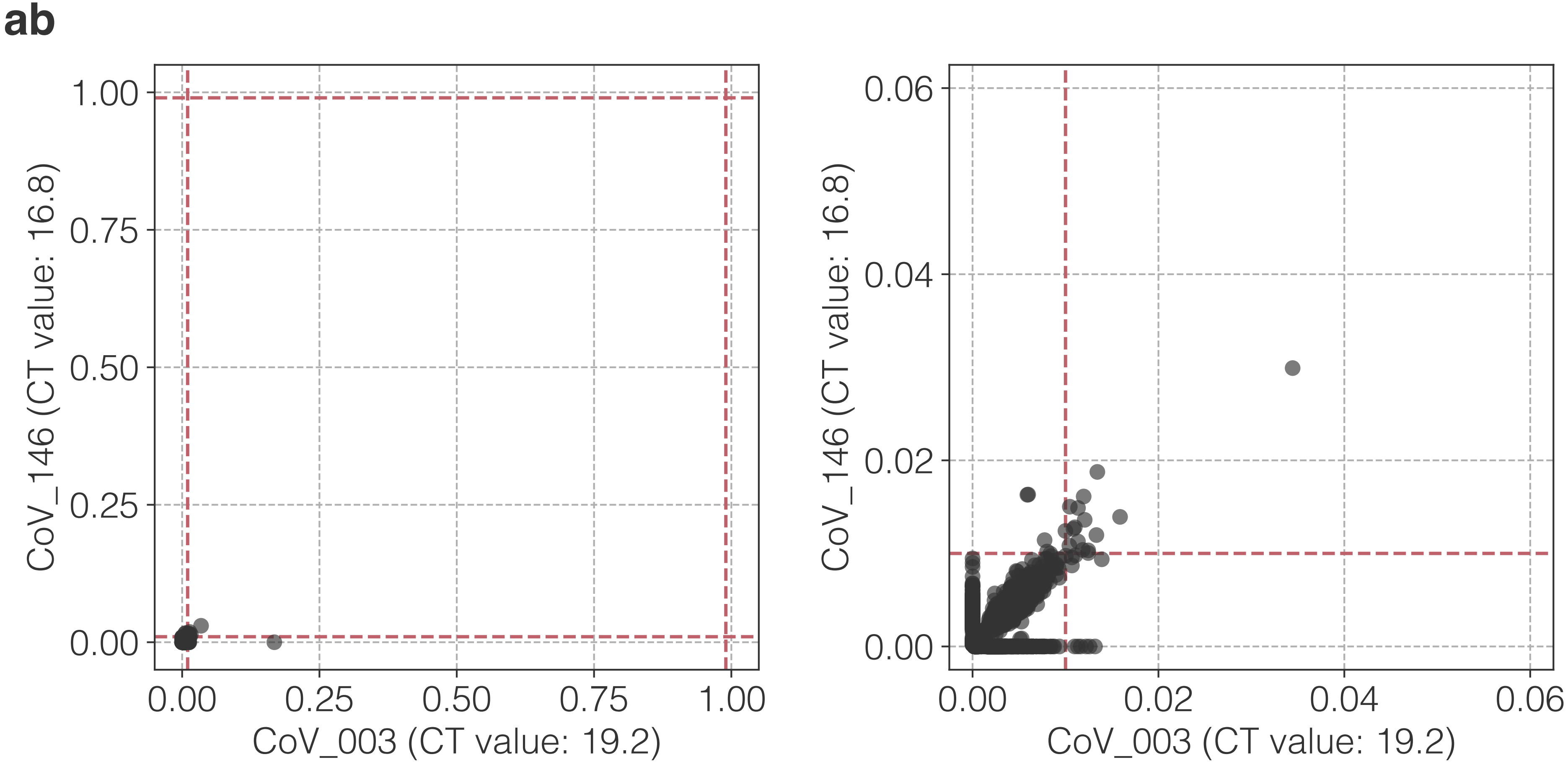

ab
CoV_146 (CT value: 16.8)
CoV_003 (CT value: 19.2)
0.00
0.25
0.50
0.75
1.00
0.00
0.02
0.04
0.06

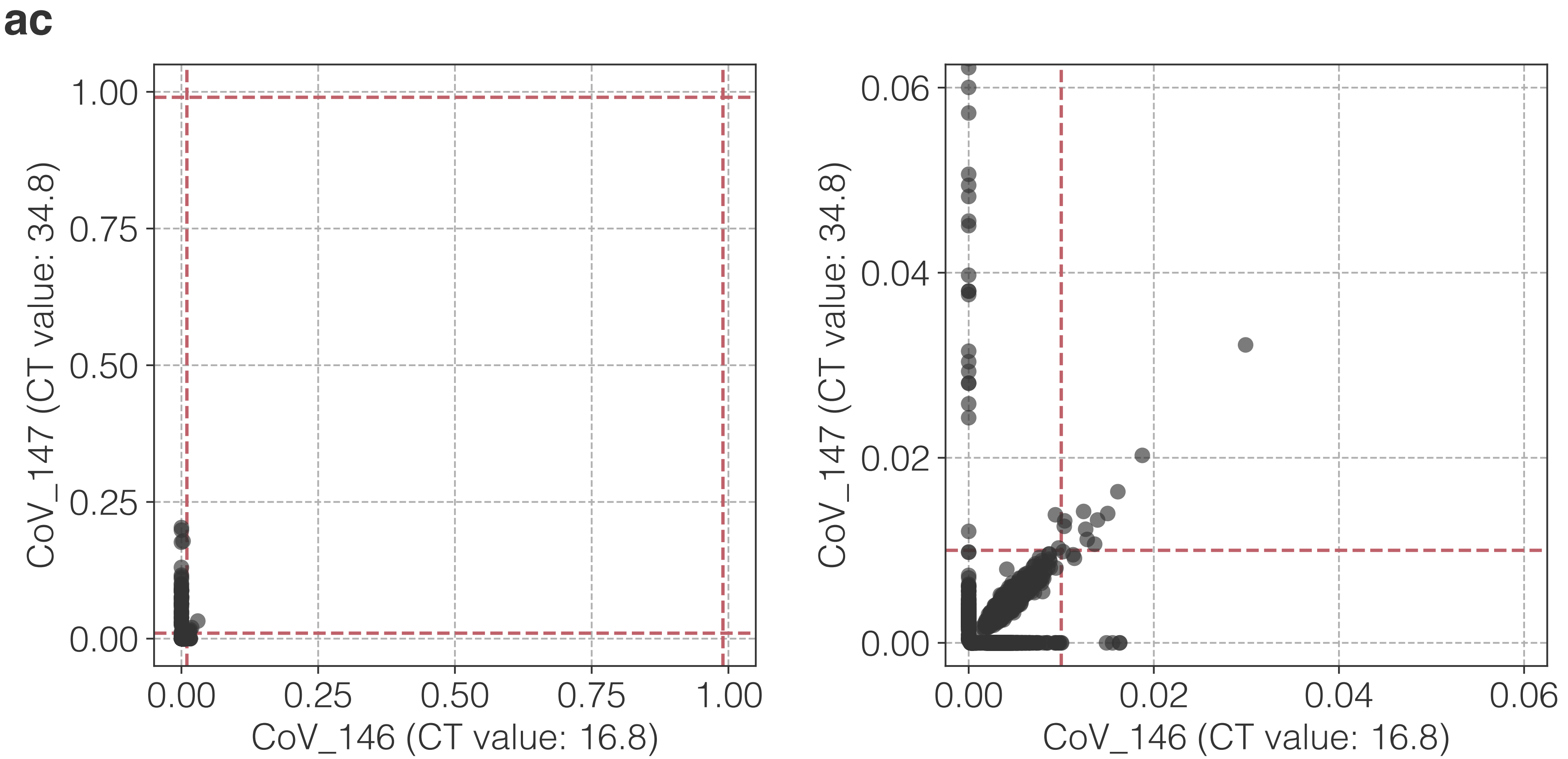
ac
1.00
0.75
0.50
0.25
0.00
CoV_147 (CT value: 34.8)
0.00
0.25
0.50
0.75
1.00
CoV_146 (CT value: 16.8)
0.06
0.04
0.02
0.00
CoV_147 (CT value: 34.8)
0.00
0.02
0.04
0.06
CoV_146 (CT value: 16.8)

**ad**

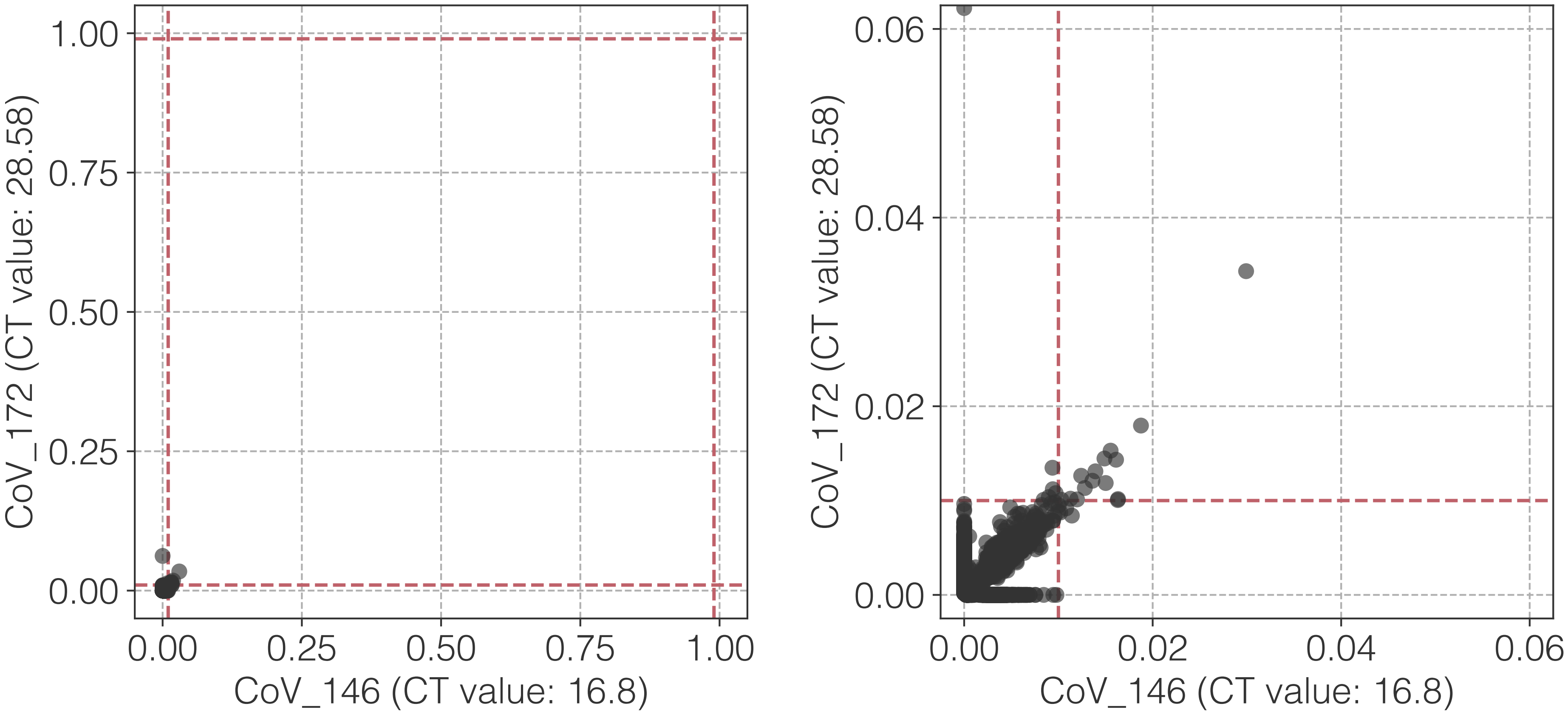

1.00
0.75
0.50
0.25
0.00
CoV_172 (CT value: 28.58)
0.00
0.25
0.50
0.75
1.00
CoV_146 (CT value: 16.8)
0.06
0.04
0.02
0.00
CoV_172 (CT value: 28.58)
0.00
0.02
0.04
0.06
CoV_146 (CT value: 16.8)

**ae**

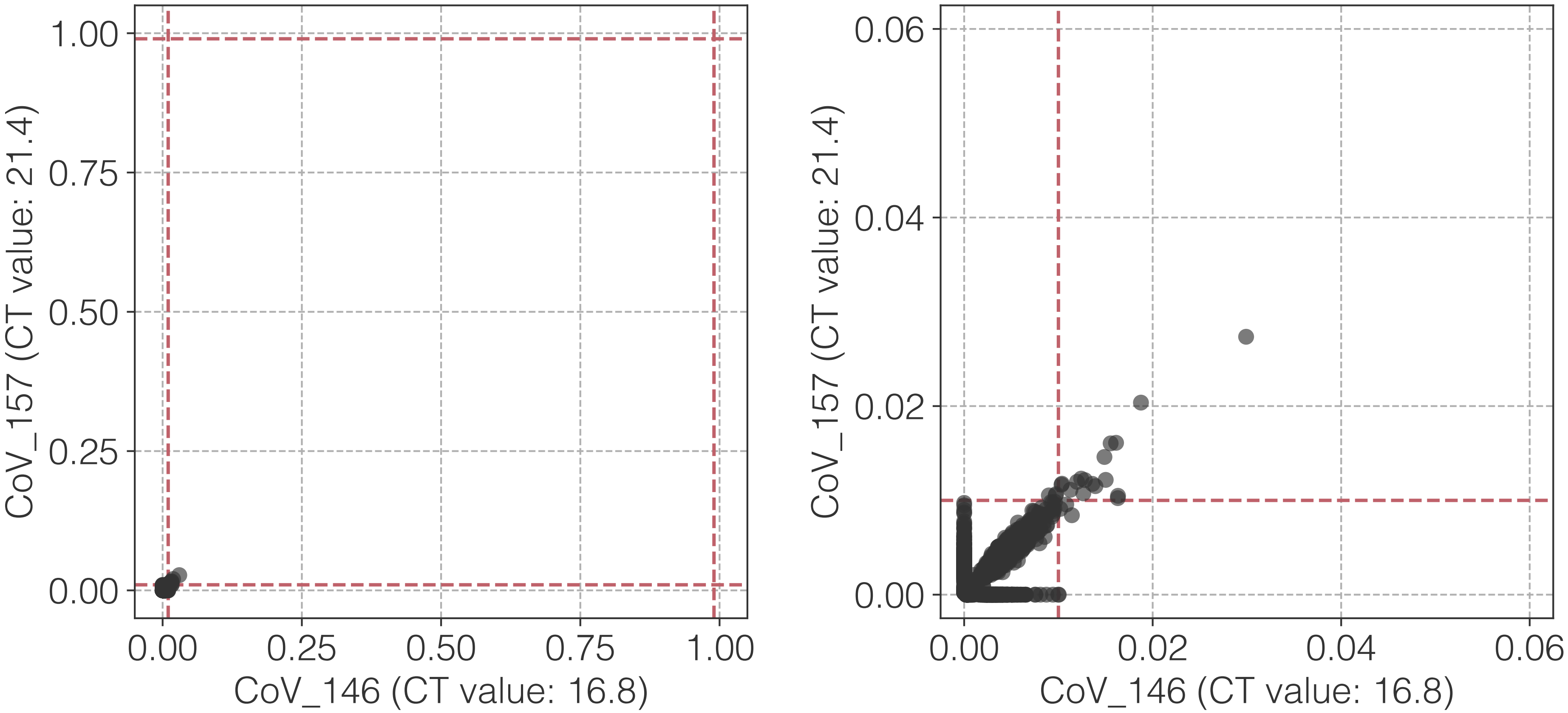

CoV_157 (CT value: 21.4)
CoV_146 (CT value: 16.8)
0.00
0.25
0.50
0.75
1.00
0.00
0.02
0.04
0.06

**af**

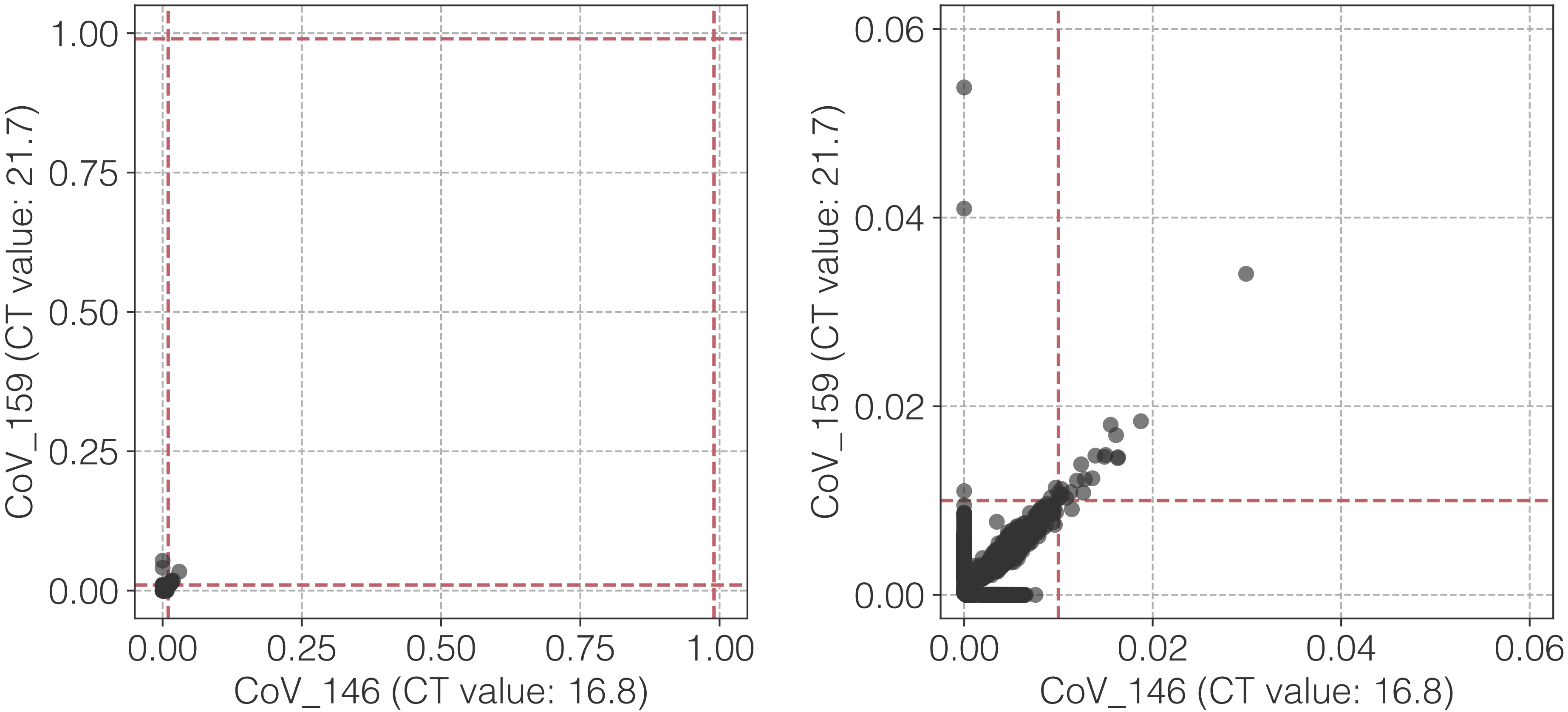

1.00
0.75
0.50
0.25
0.00
CoV_159 (CT value: 21.7)
0.00
0.25
0.50
0.75
1.00
CoV_146 (CT value: 16.8)
0.06
0.04
0.02
0.00
CoV_159 (CT value: 21.7)
0.00
0.02
0.04
0.06
CoV_146 (CT value: 16.8)

ag

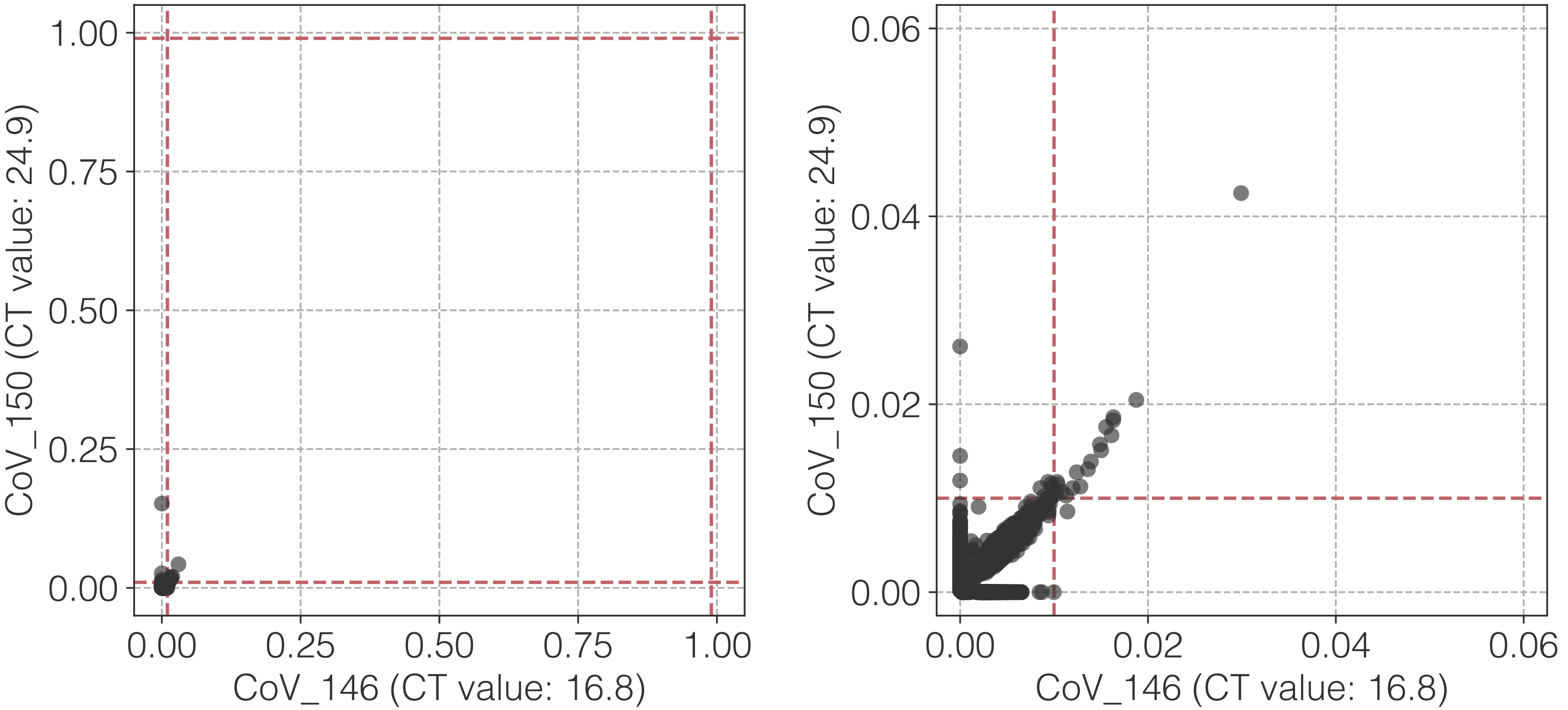

1.00
0.75
0.50
0.25
0.00
CoV_150 (CT value: 24.9)
0.00
0.25
0.50
0.75
1.00
CoV_146 (CT value: 16.8)
0.06
0.04
0.02
0.00
CoV_150 (CT value: 24.9)
0.00
0.02
0.04
0.06
CoV_146 (CT value: 16.8)

**ah**

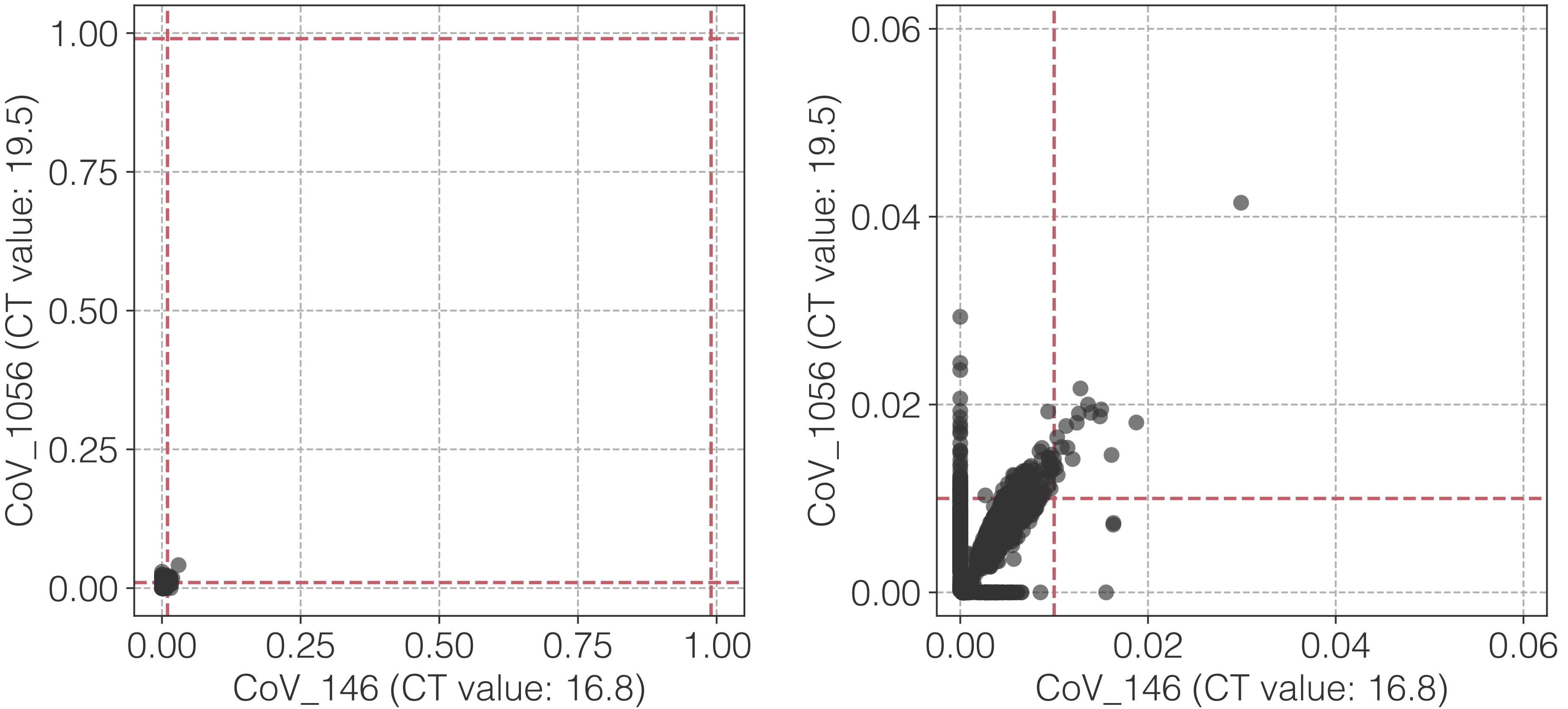

CoV_1056 (CT value: 19.5)
CoV_146 (CT value: 16.8)
0.00
0.25
0.50
0.75
1.00
0.02
0.04
0.06

**ai**

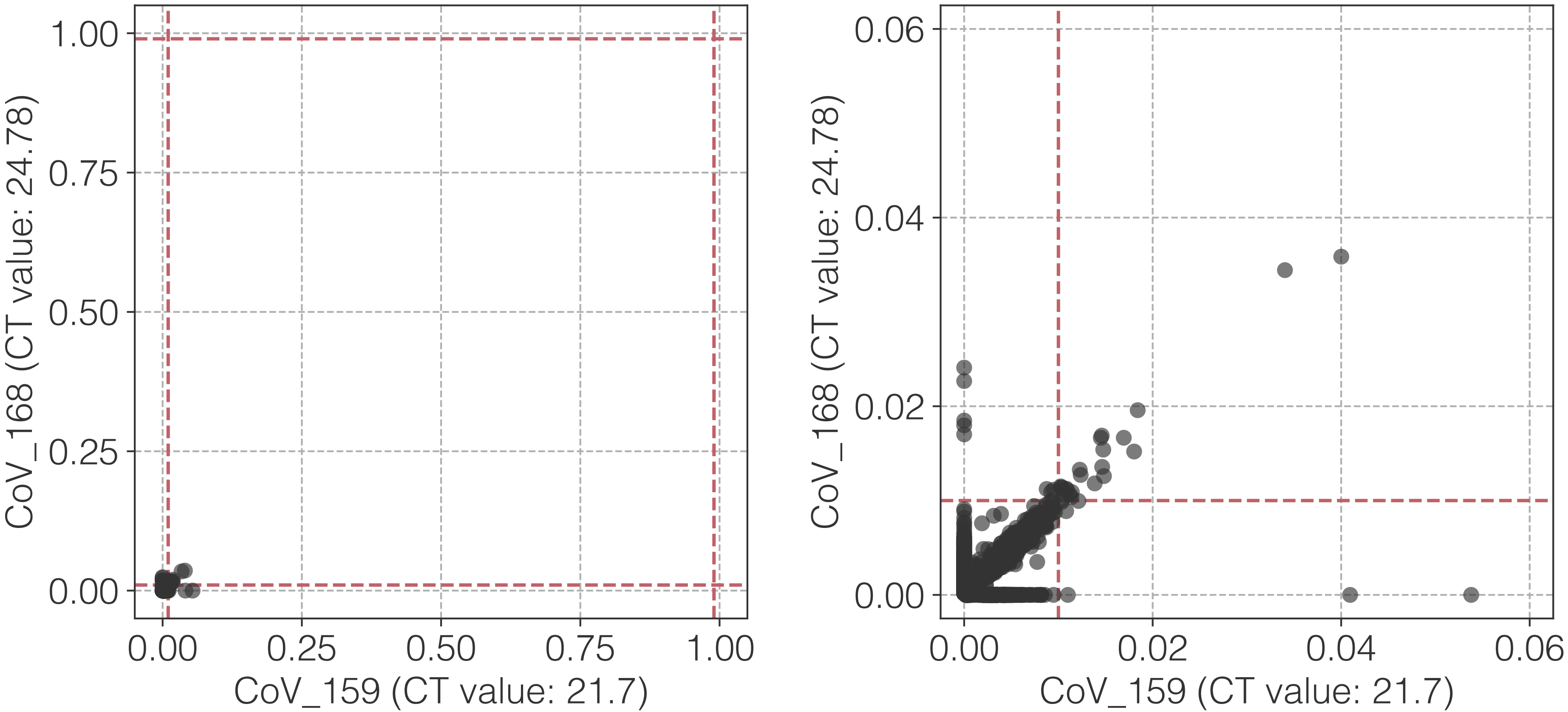

CoV_168 (CT value: 24.78)
CoV_159 (CT value: 21.7)
0.00
0.25
0.50
0.75
1.00
CoV_168 (CT value: 24.78)
CoV_159 (CT value: 21.7)
0.00
0.02
0.04
0.06

**aj**

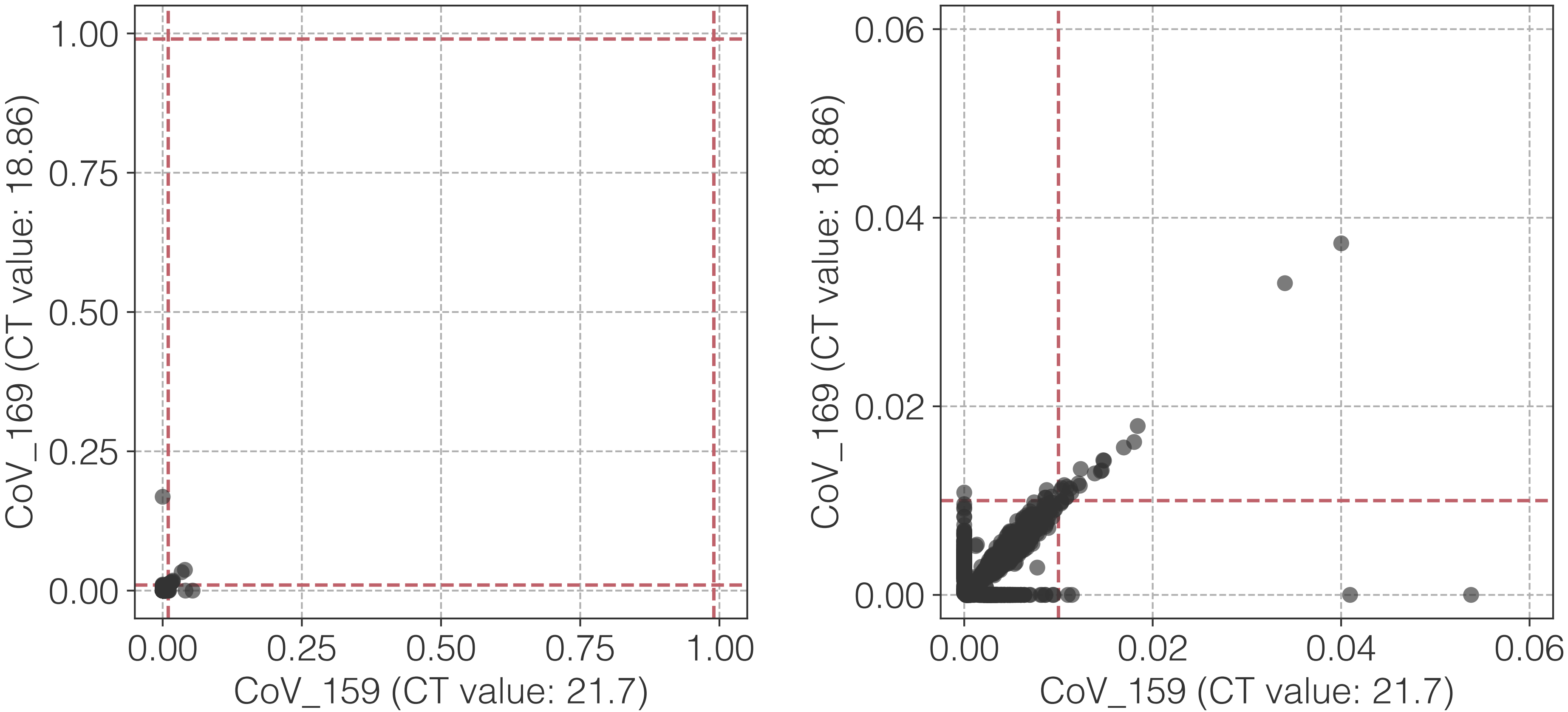
CoV_169 (CT value: 18.86)
CoV_159 (CT value: 21.7)
1.00
0.75
0.50
0.25
0.00
0.00
0.25
0.50
0.75
1.00
CoV_169 (CT value: 18.86)
CoV_159 (CT value: 21.7)
0.06
0.04
0.02
0.00
0.00
0.02
0.04
0.06

**ak**

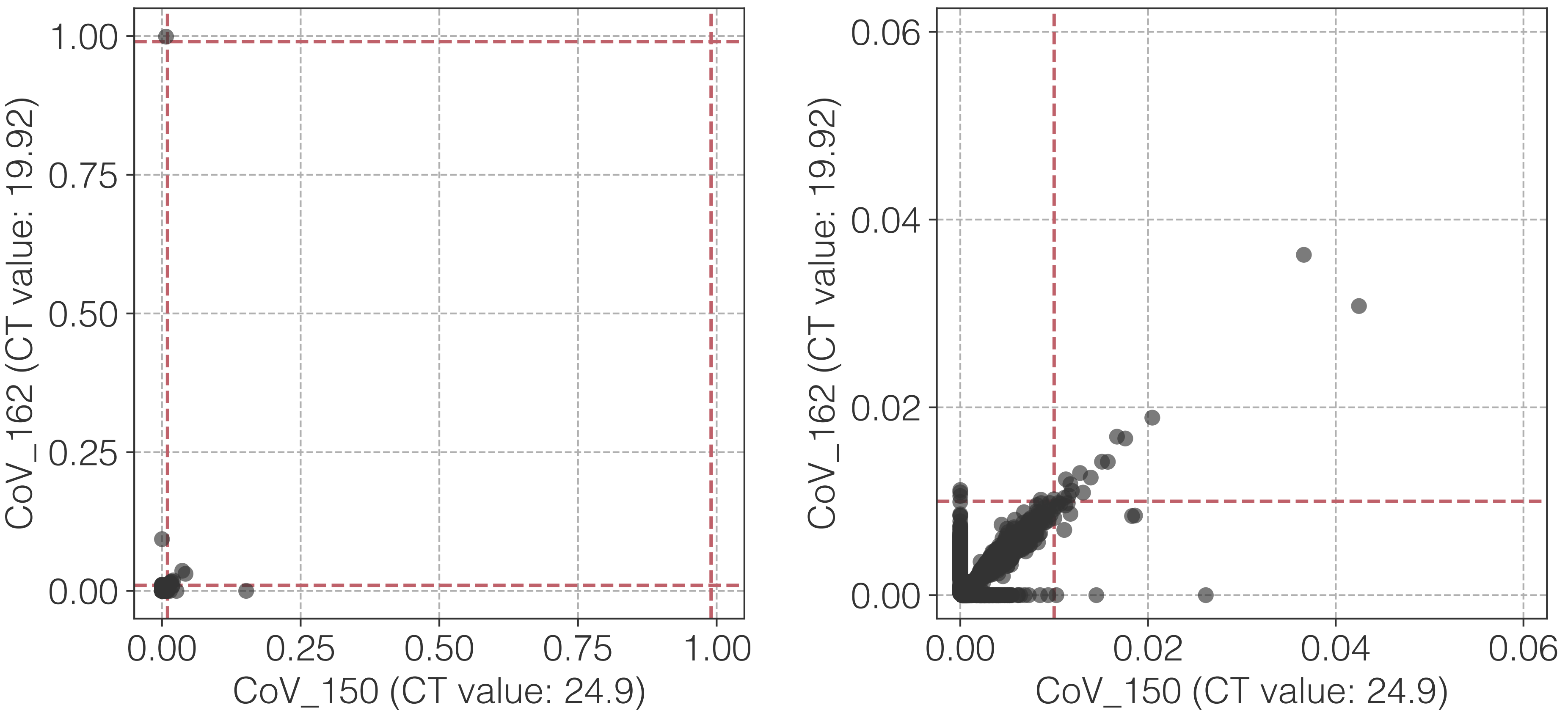

CoV_162 (CT value: 19.92)
CoV_150 (CT value: 24.9)
1.00
0.75
0.50
0.25
0.00
0.00
0.25
0.50
0.75
1.00
CoV_162 (CT value: 19.92)
CoV_150 (CT value: 24.9)
0.06
0.04
0.02
0.00
0.00
0.02
0.04
0.06

al

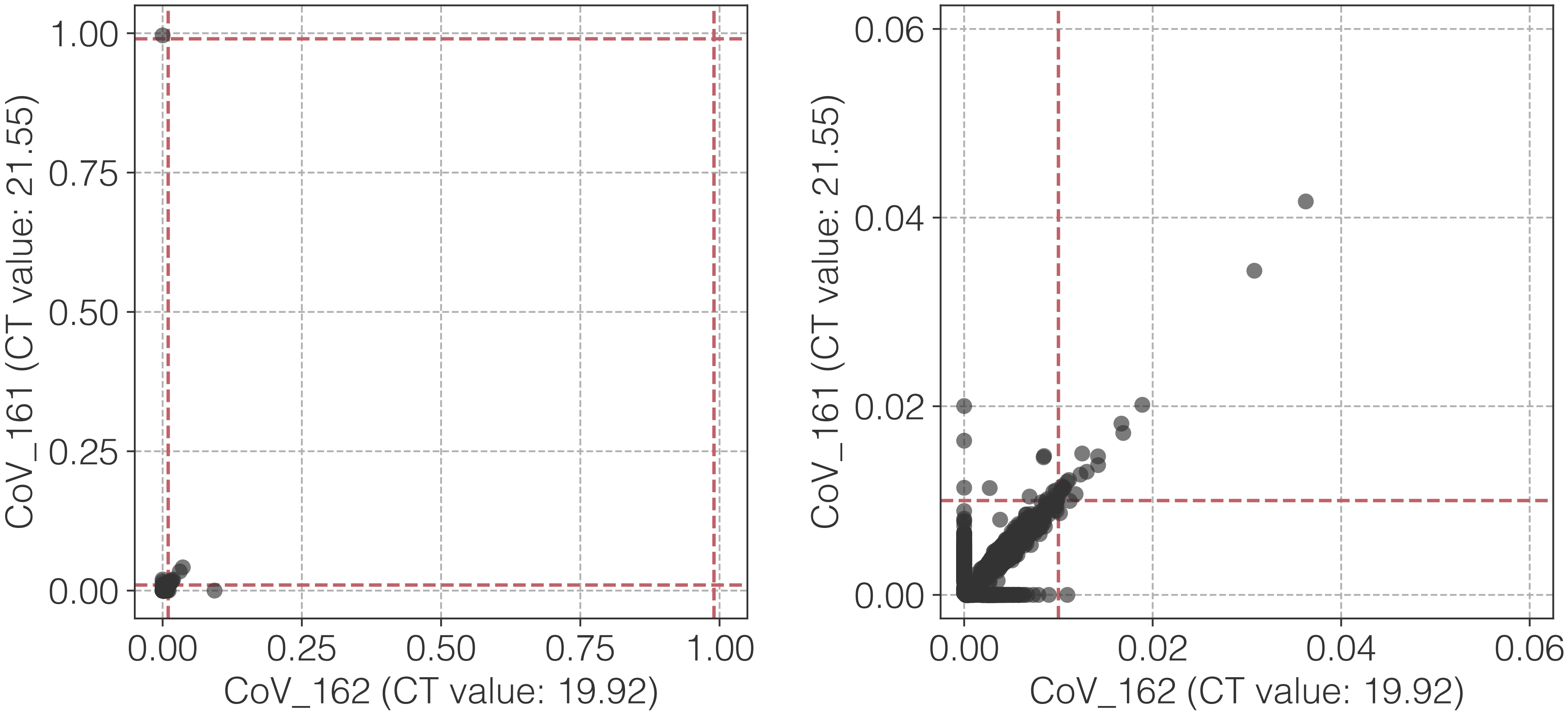

1.00
0.75
0.50
0.25
0.00
CoV_161 (CT value: 21.55)
0.00
0.25
0.50
0.75
1.00
CoV_162 (CT value: 19.92)
0.06
0.04
0.02
0.00
CoV_161 (CT value: 21.55)
0.00
0.02
0.04
0.06
CoV_162 (CT value: 19.92)

**am**

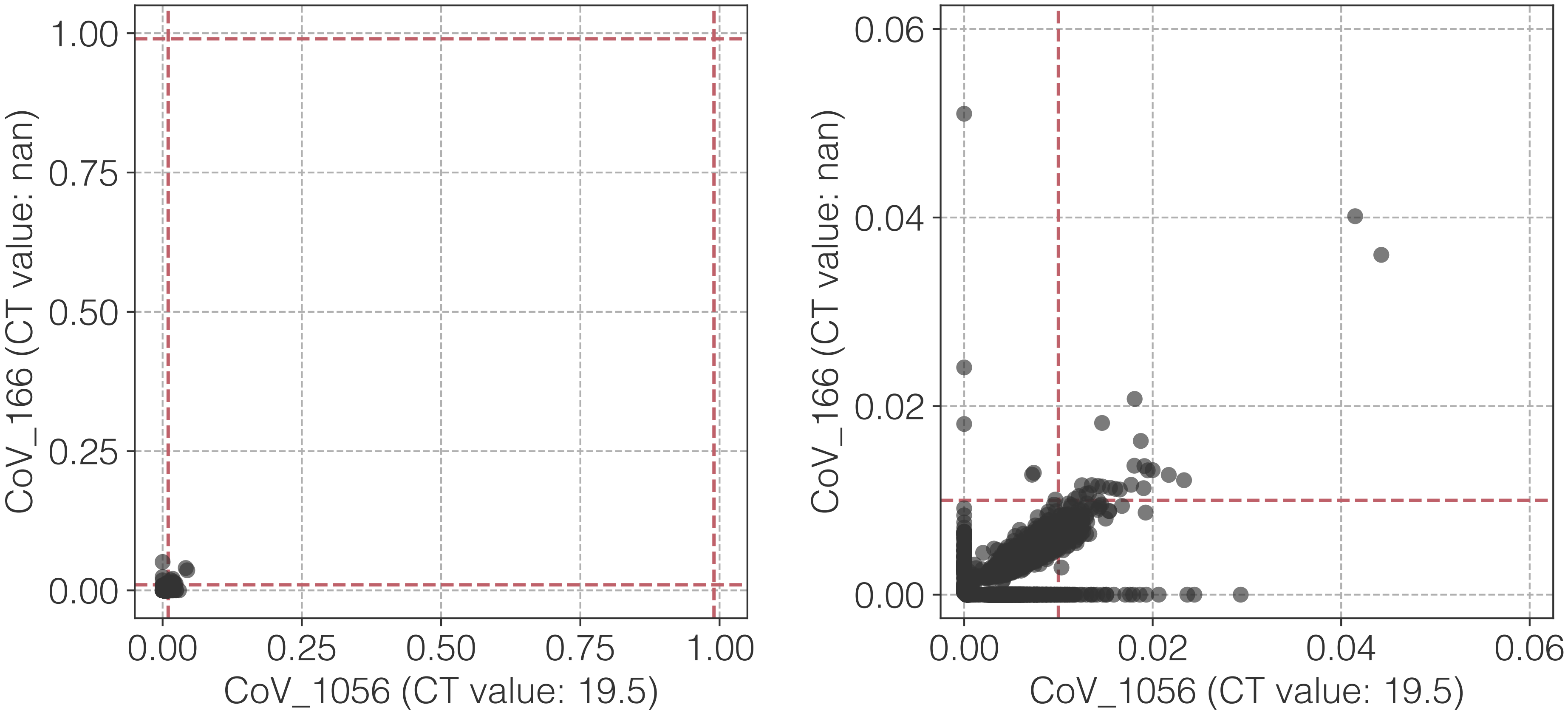

CoV_166 (CT value: nan)
1.00
0.75
0.50
0.25
0.00
0.00
0.25
0.50
0.75
1.00
CoV_1056 (CT value: 19.5)
CoV_166 (CT value: nan)
0.06
0.04
0.02
0.00
0.00
0.02
0.04
0.06
CoV_1056 (CT value: 19.5)

**an**

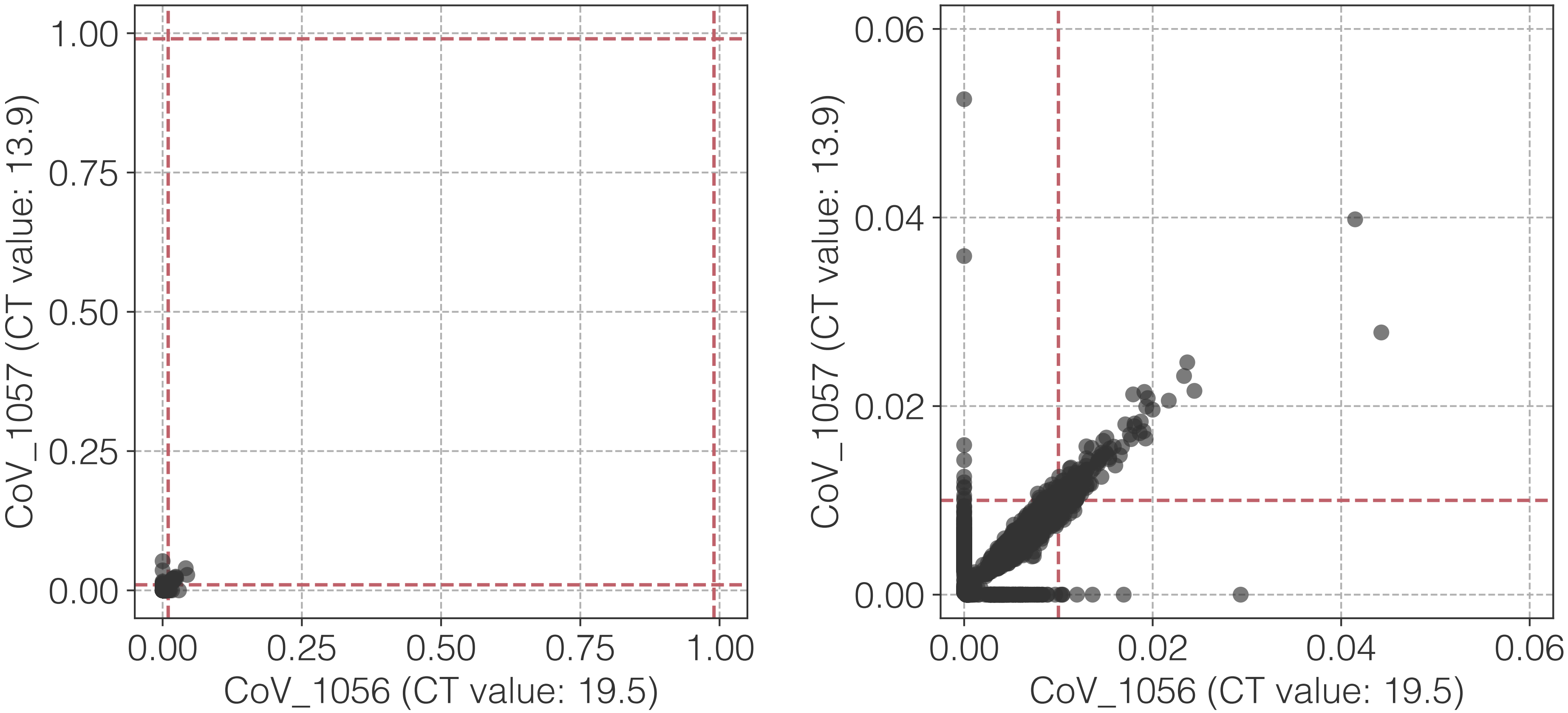
1.00
0.75
0.50
0.25
0.00
CoV_1057 (CT value: 13.9)
0.00
0.25
0.50
0.75
1.00
CoV_1056 (CT value: 19.5)
0.06
0.04
0.02
0.00
CoV_1057 (CT value: 13.9)
0.00
0.02
0.04
0.06
CoV_1056 (CT value: 19.5)

**ao**

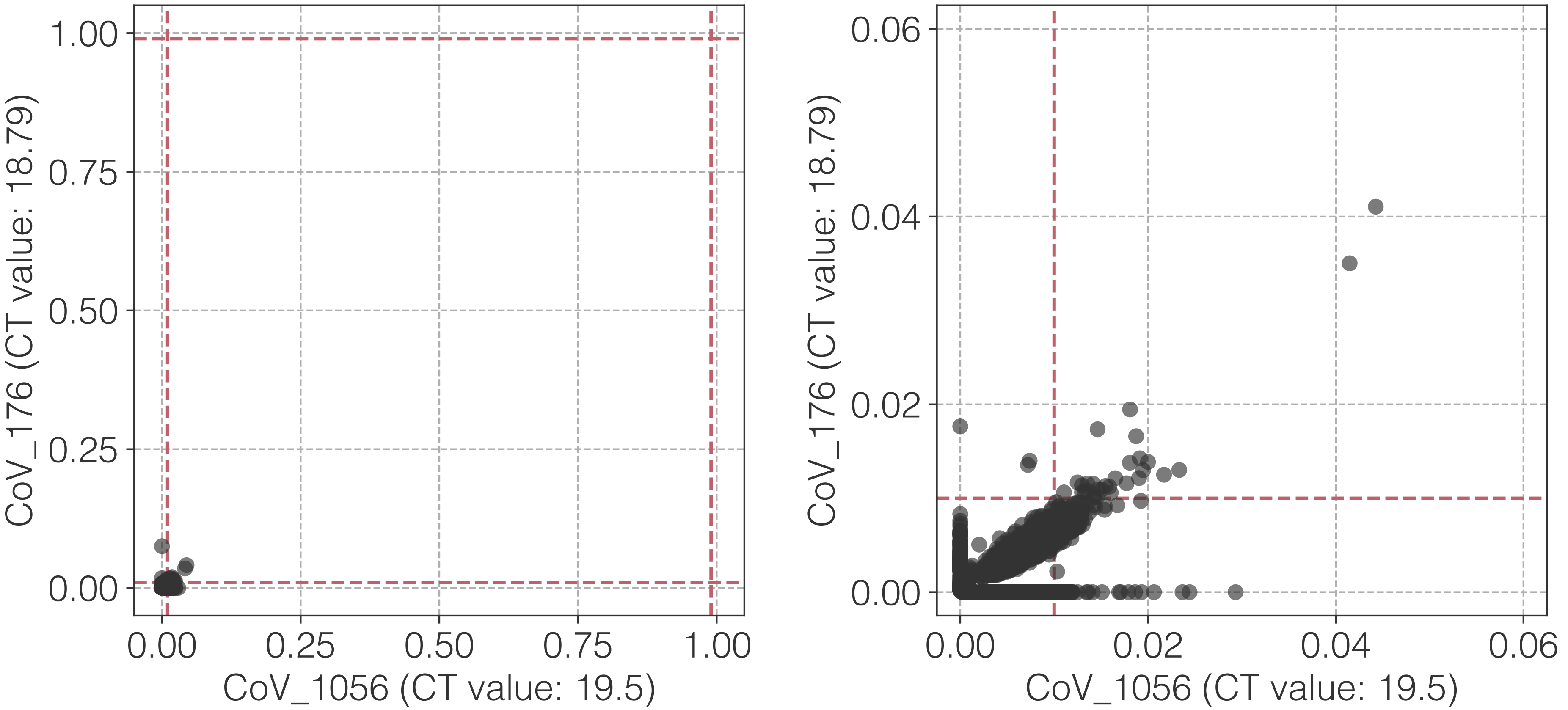

CoV_176 (CT value: 18.79)
CoV_1056 (CT value: 19.5)
0.00
0.25
0.50
0.75
1.00
CoV_176 (CT value: 18.79)
CoV_1056 (CT value: 19.5)
0.00
0.02
0.04
0.06

**ap**

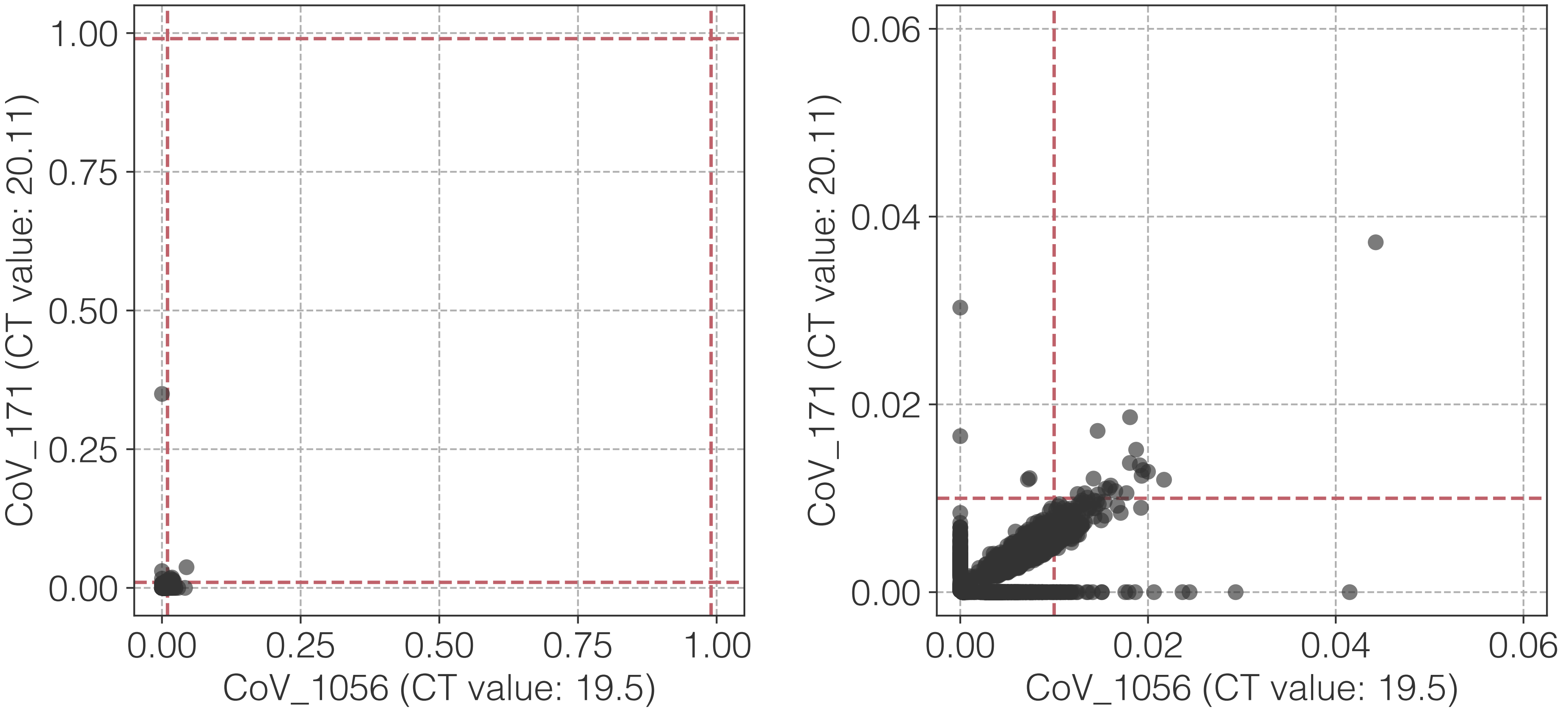

1.00
0.75
0.50
0.25
0.00
0.00
0.25
0.50
0.75
1.00
CoV_171 (CT value: 20.11)
CoV_1056 (CT value: 19.5)
0.06
0.04
0.02
0.00
0.00
0.02
0.04
0.06
CoV_171 (CT value: 20.11)
CoV_1056 (CT value: 19.5)

**aq**

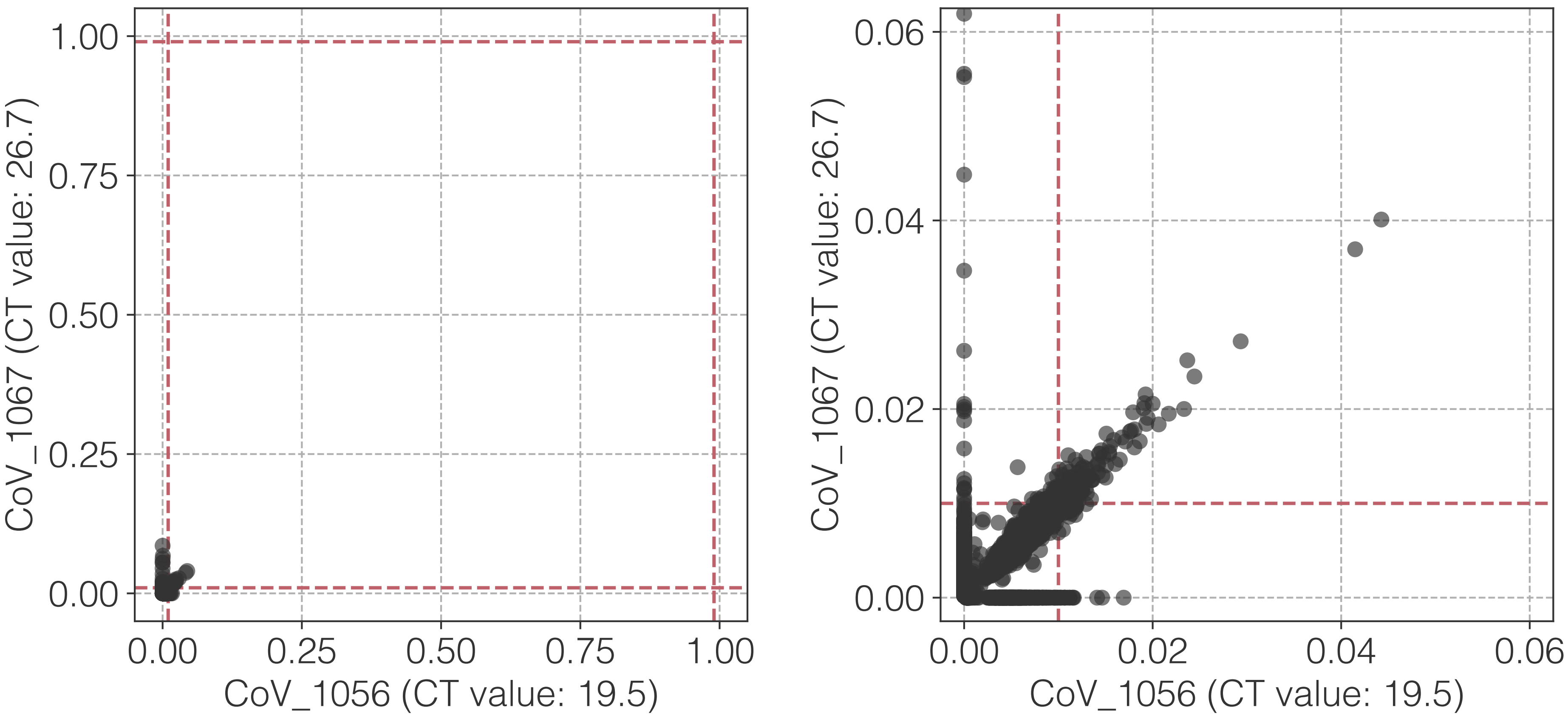

1.00
0.75
0.50
0.25
0.00
CoV_1067 (CT value: 26.7)
0.00
0.25
0.50
0.75
1.00
CoV_1056 (CT value: 19.5)
0.06
0.04
0.02
0.00
CoV_1067 (CT value: 26.7)
0.00
0.02
0.04
0.06
CoV_1056 (CT value: 19.5)

**ar**

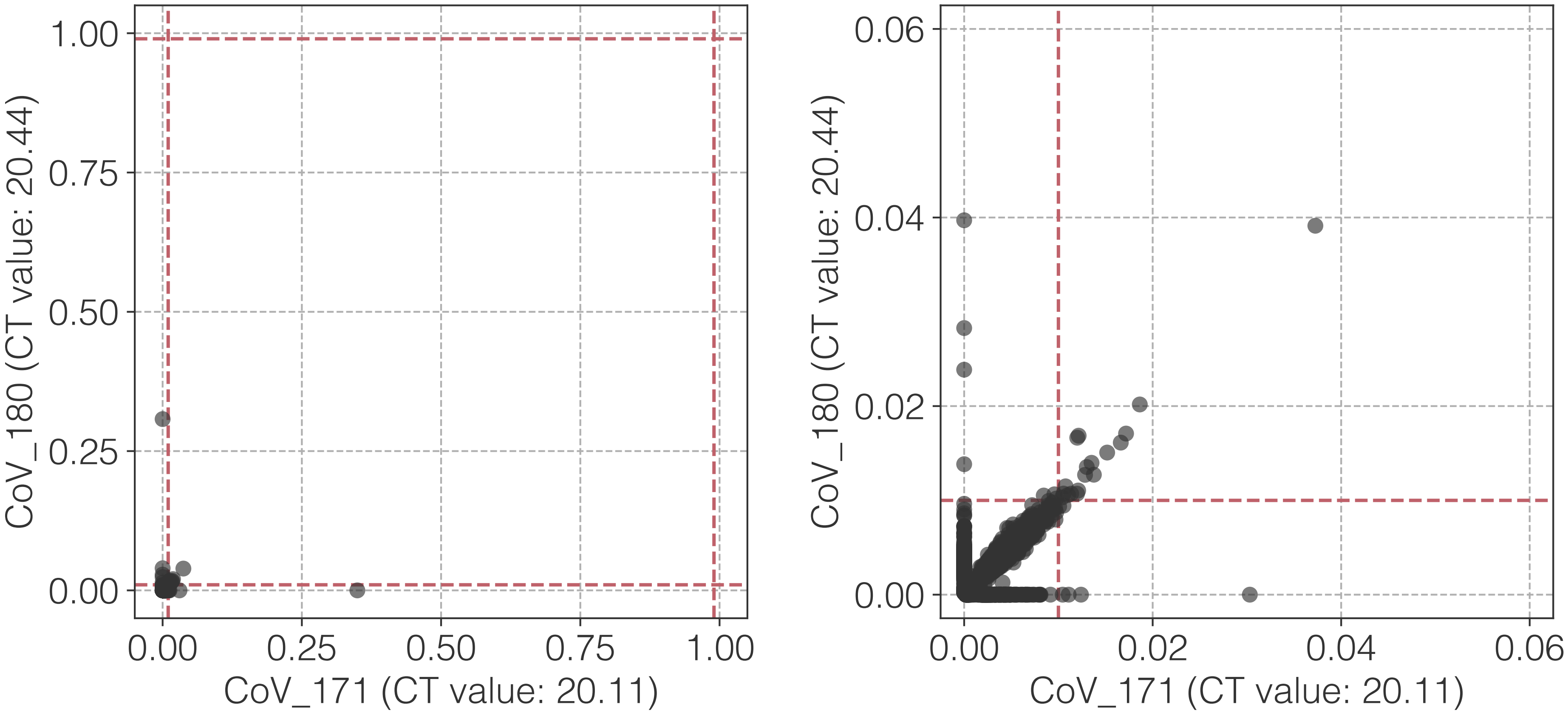

1.00
0.75
0.50
0.25
0.00
CoV_180 (CT value: 20.44)
0.00
0.25
0.50
0.75
1.00
CoV_171 (CT value: 20.11)
0.06
0.04
0.02
0.00
CoV_180 (CT value: 20.44)
0.00
0.02
0.04
0.06
CoV_171 (CT value: 20.11)

**as**

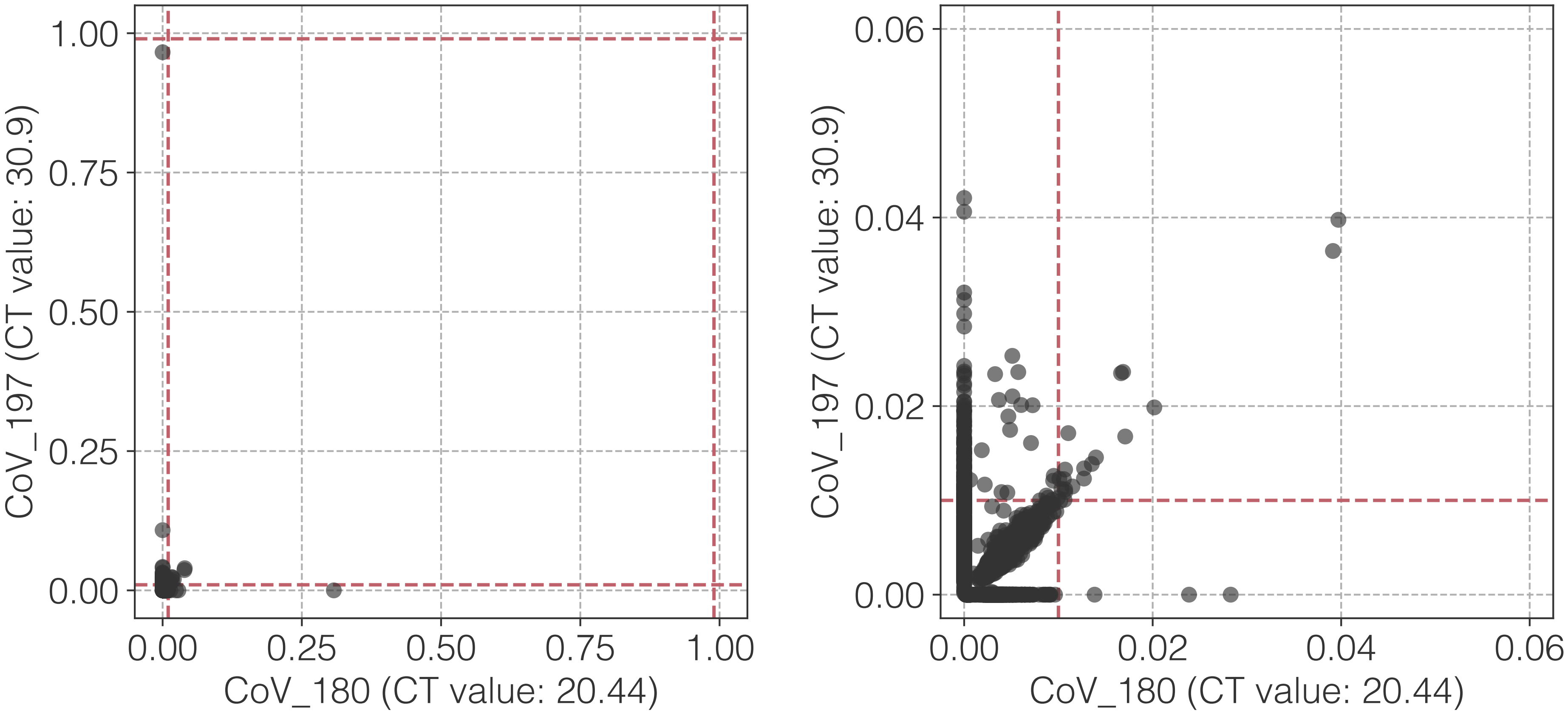
1.00
0.75
0.50
0.25
0.00
CoV_197 (CT value: 30.9)
0.00
0.25
0.50
0.75
1.00
CoV_180 (CT value: 20.44)
0.06
0.04
0.02
0.00
CoV_197 (CT value: 30.9)
0.00
0.02
0.04
0.06
CoV_180 (CT value: 20.44)

**at**

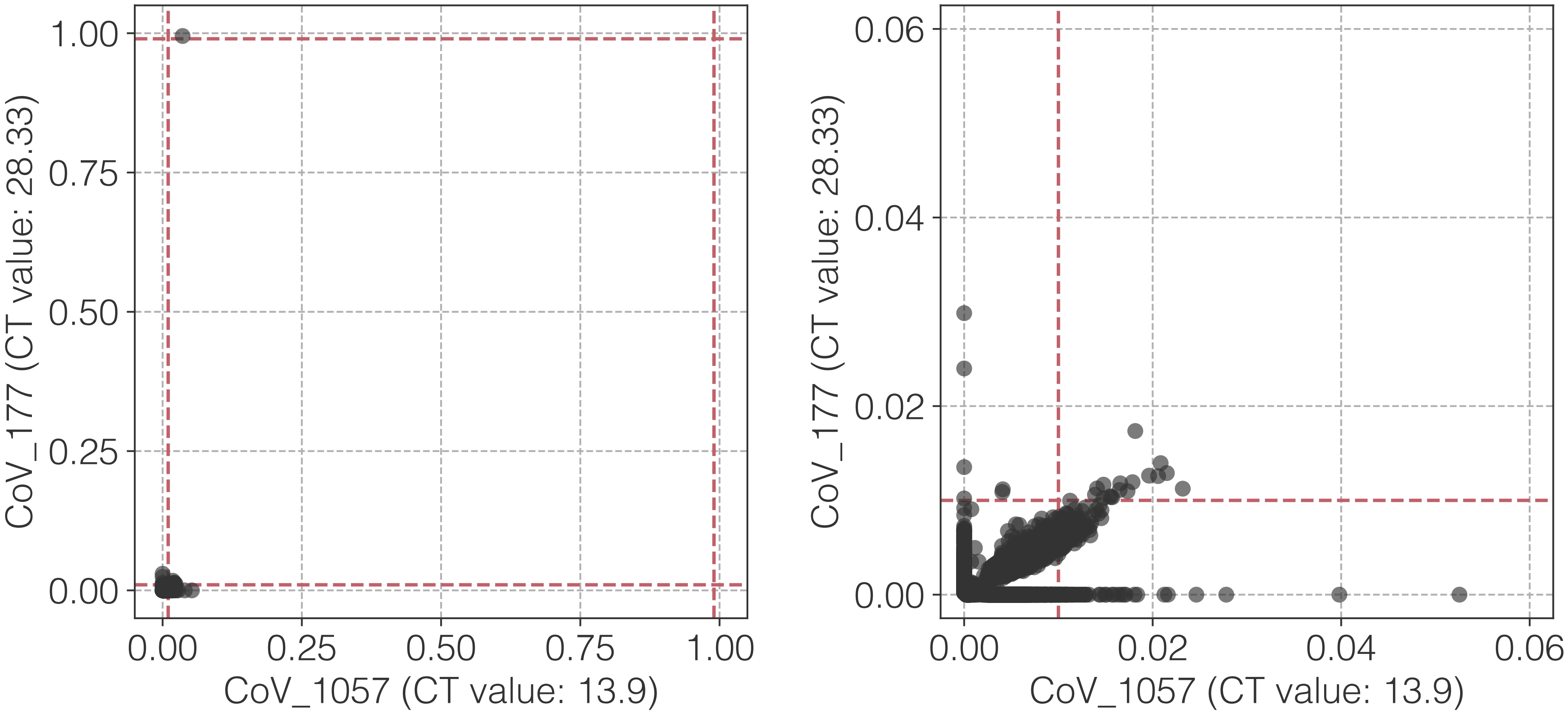

1.00
0.75
0.50
0.25
0.00
CoV_177 (CT value: 28.33)
0.00
0.25
0.50
0.75
1.00
CoV_1057 (CT value: 13.9)
0.06
0.04
0.02
0.00
CoV_177 (CT value: 28.33)
0.00
0.02
0.04
0.06
CoV_1057 (CT value: 13.9)

**au**

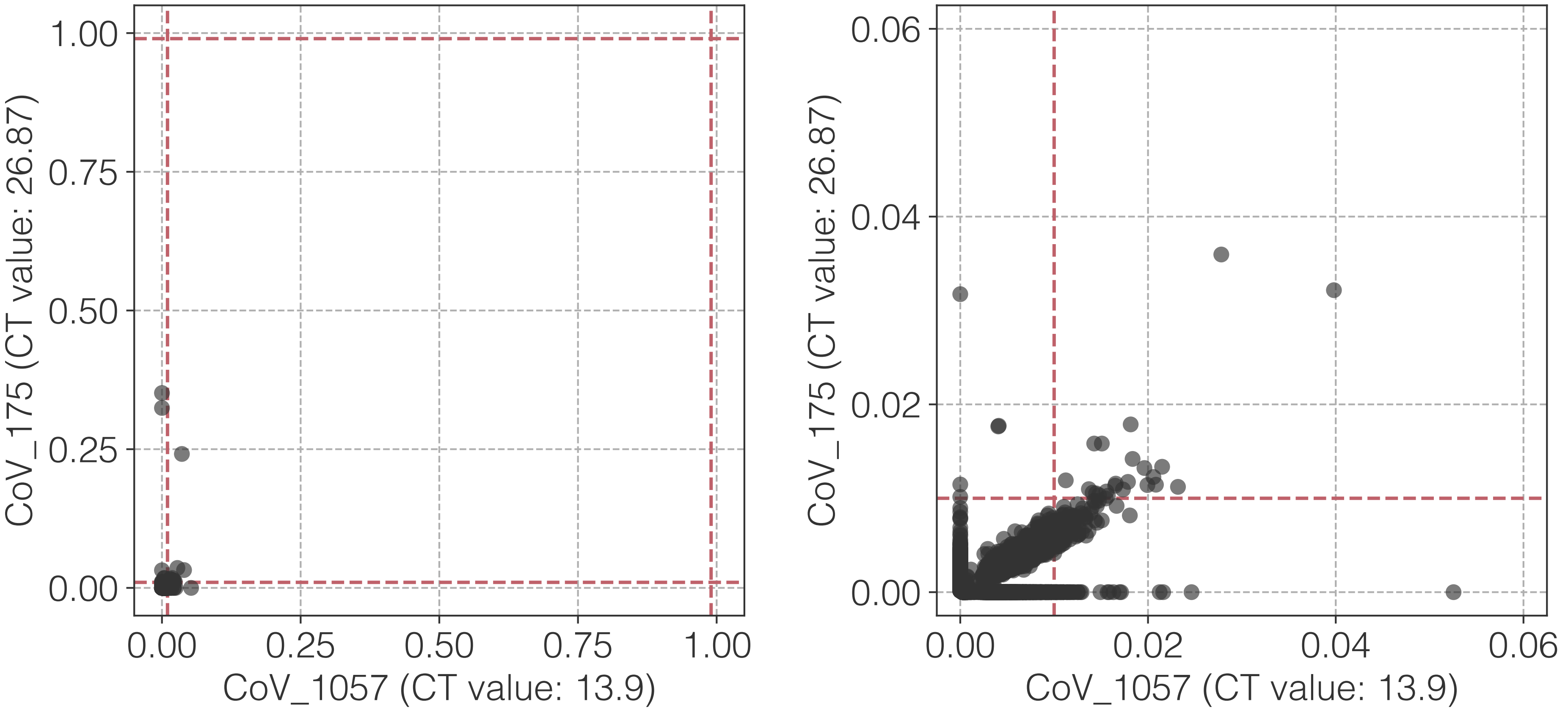

1.00
0.75
0.50
0.25
0.00
CoV_175 (CT value: 26.87)
0.00
0.25
0.50
0.75
1.00
CoV_1057 (CT value: 13.9)
0.06
0.04
0.02
0.00
CoV_175 (CT value: 26.87)
0.00
0.02
0.04
0.06
CoV_1057 (CT value: 13.9)

**av**

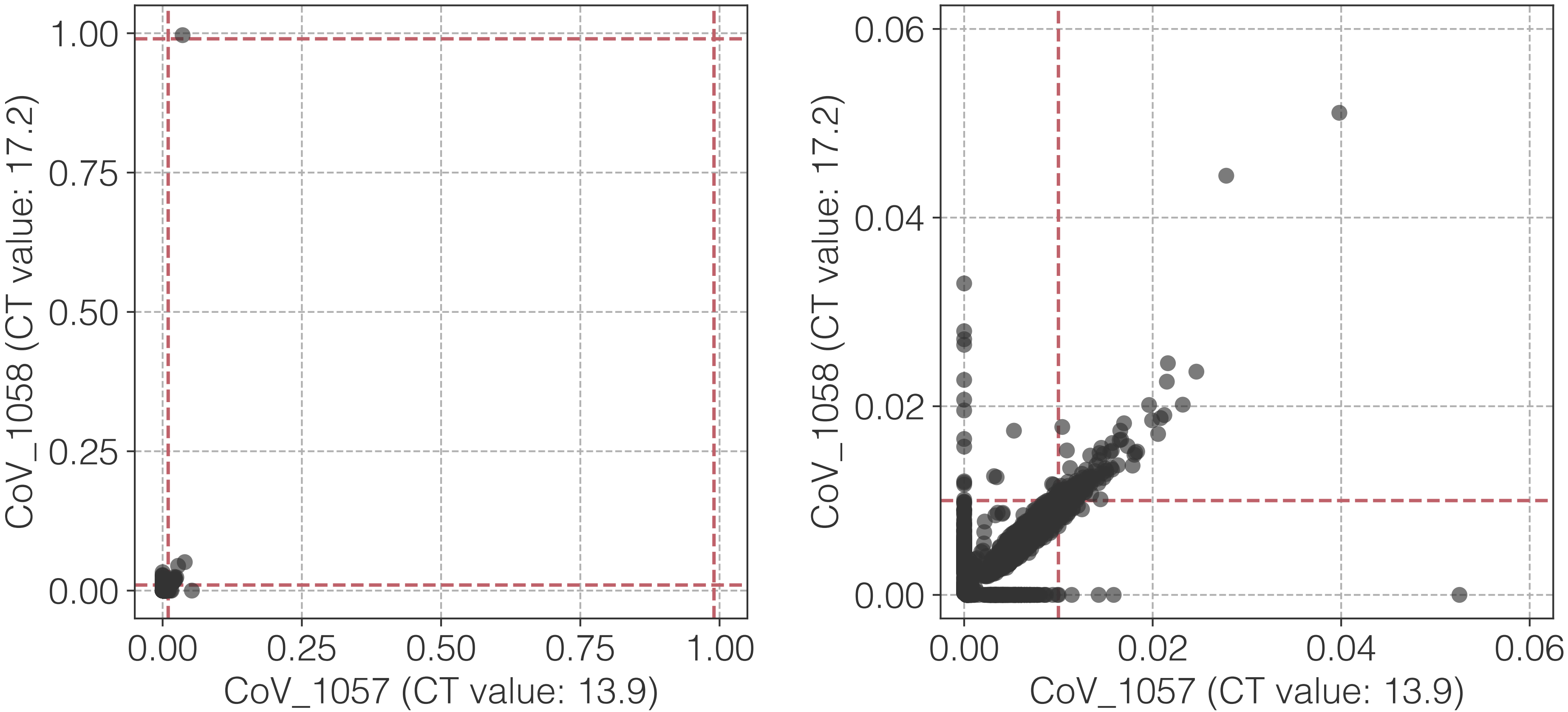

1.00
0.75
0.50
0.25
0.00
CoV_1058 (CT value: 17.2)
0.00
0.25
0.50
0.75
1.00
CoV_1057 (CT value: 13.9)
0.06
0.04
0.02
0.00
CoV_1058 (CT value: 17.2)
0.00
0.02
0.04
0.06
CoV_1057 (CT value: 13.9)

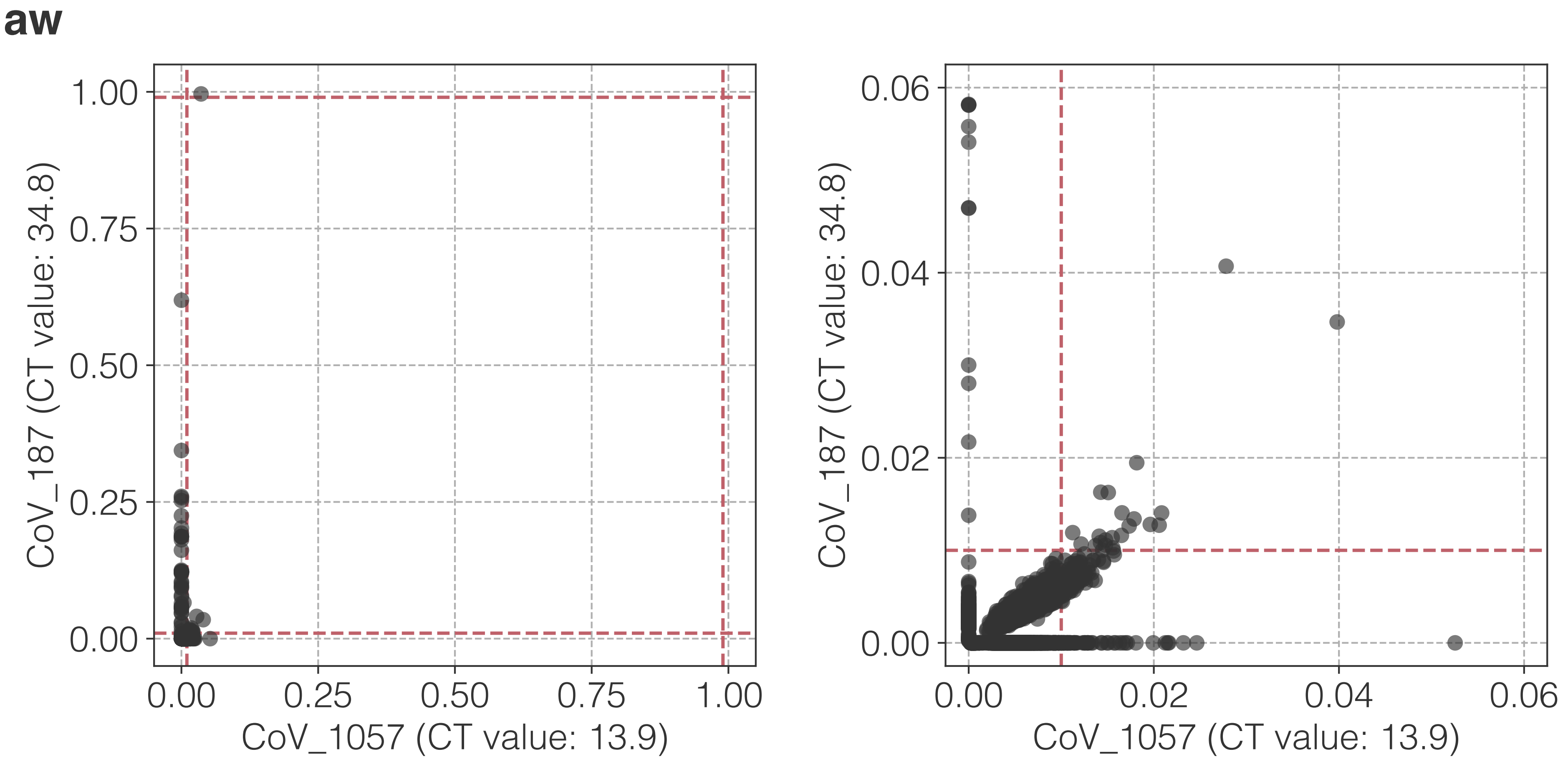
aw
CoV_187 (CT value: 34.8)
CoV_1057 (CT value: 13.9)
0.00
0.25
0.50
0.75
1.00
0.02
0.04
0.06

**ax**

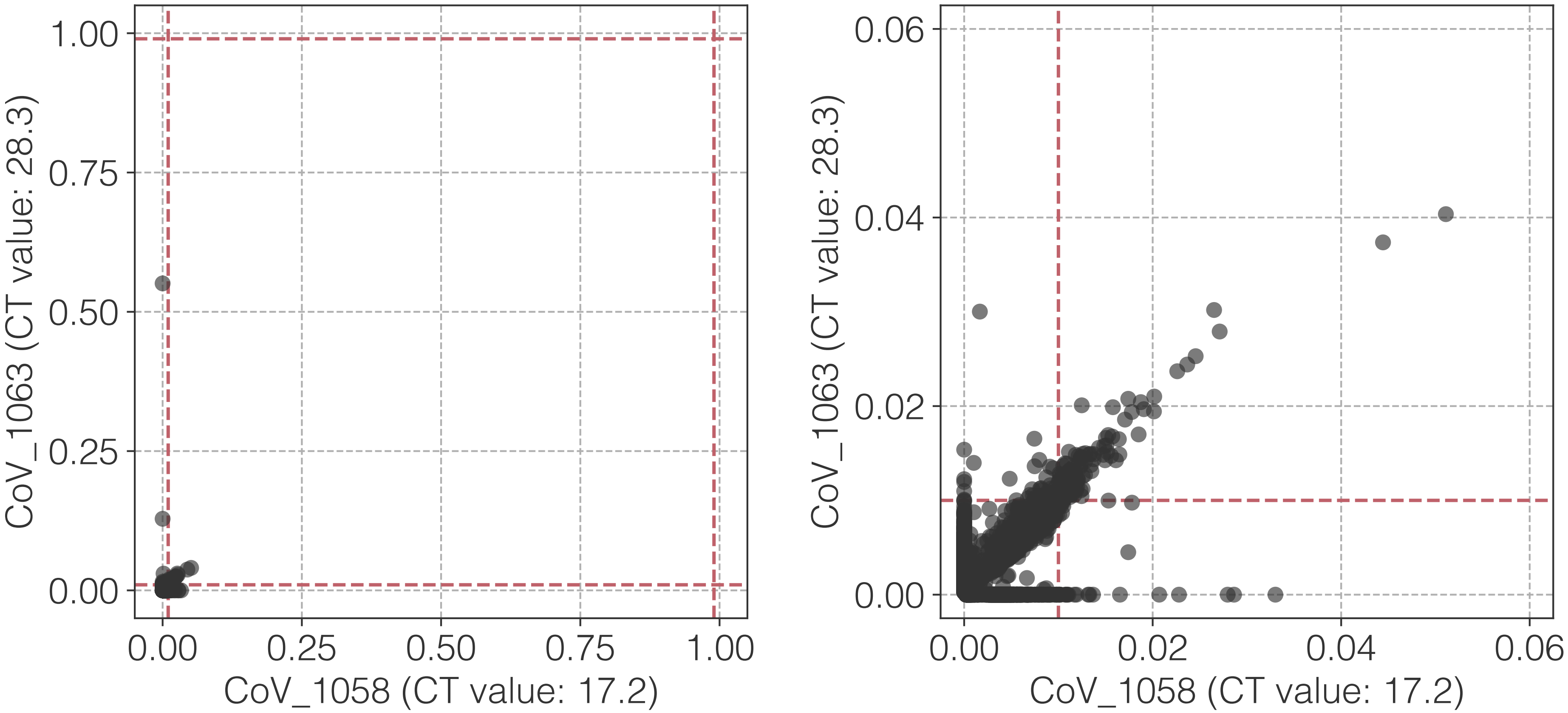

1.00
0.75
0.50
0.25
0.00
CoV_1063 (CT value: 28.3)
0.00
0.25
0.50
0.75
1.00
CoV_1058 (CT value: 17.2)
0.06
0.04
0.02
0.00
CoV_1063 (CT value: 28.3)
0.00
0.02
0.04
0.06
CoV_1058 (CT value: 17.2)

**ay**

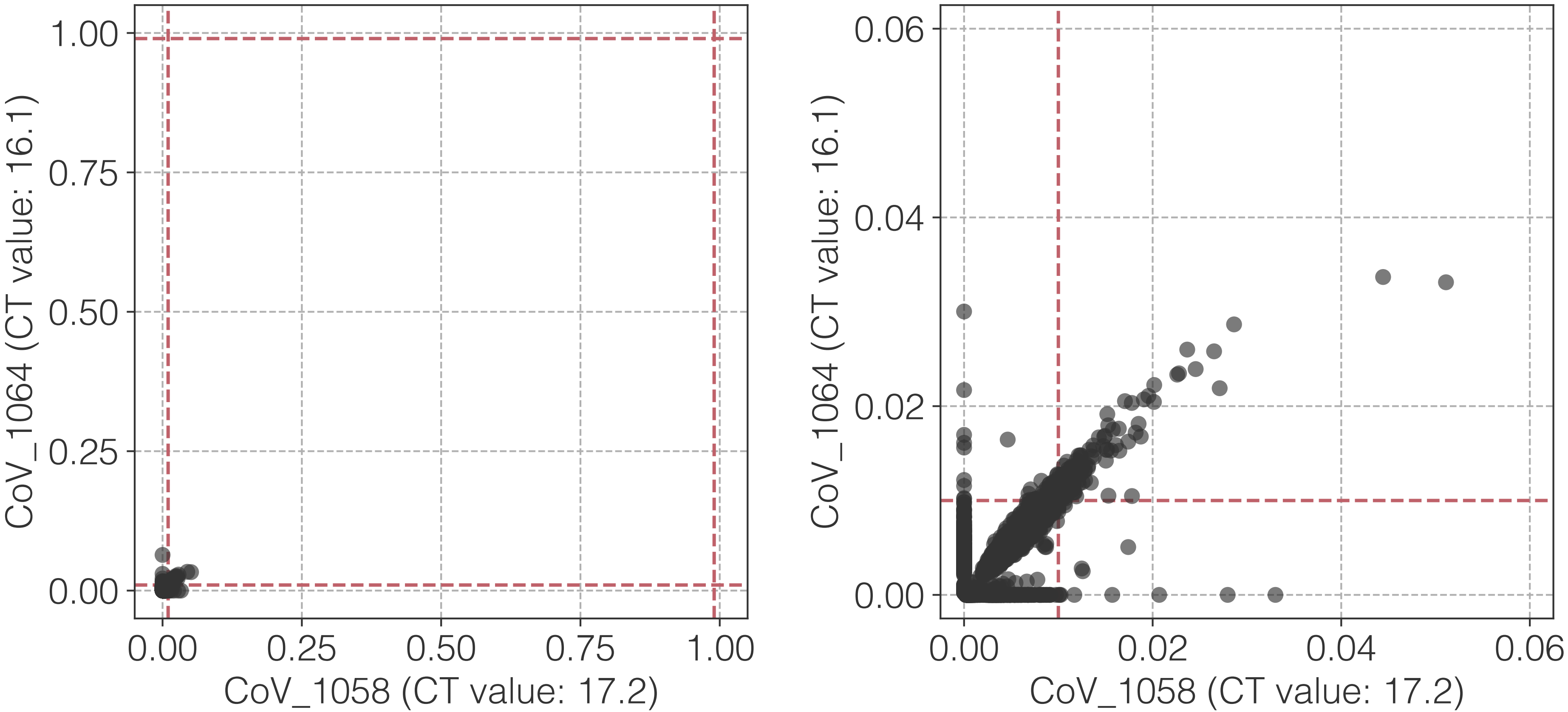

CoV_1064 (CT value: 16.1)
CoV_1058 (CT value: 17.2)
0.00
0.25
0.50
0.75
1.00
CoV_1064 (CT value: 16.1)
CoV_1058 (CT value: 17.2)
0.00
0.02
0.04
0.06

**az**

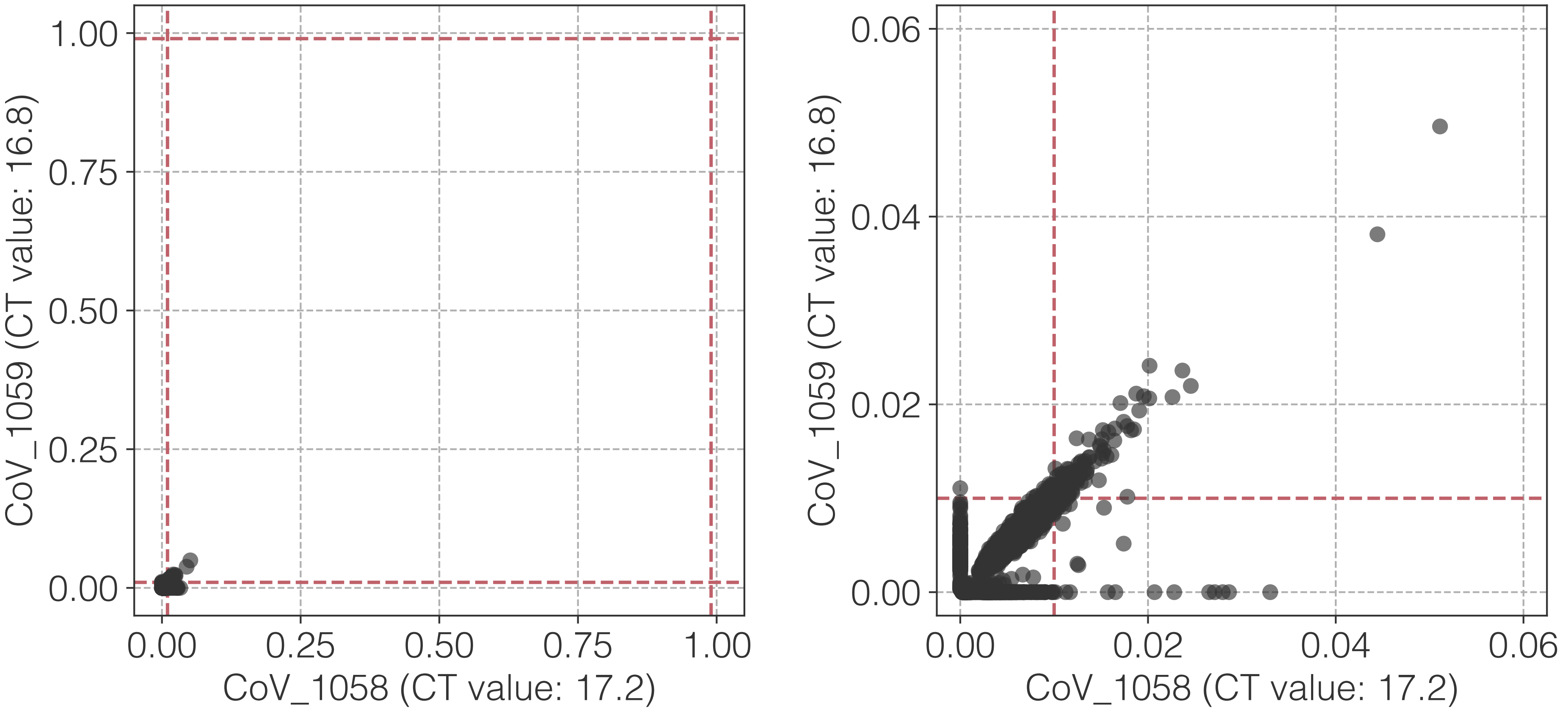

1.00
0.75
0.50
0.25
0.00
CoV_1059 (CT value: 16.8)
0.00
0.25
0.50
0.75
1.00
CoV_1058 (CT value: 17.2)
0.06
0.04
0.02
0.00
CoV_1059 (CT value: 16.8)
0.00
0.02
0.04
0.06
CoV_1058 (CT value: 17.2)

**ba**

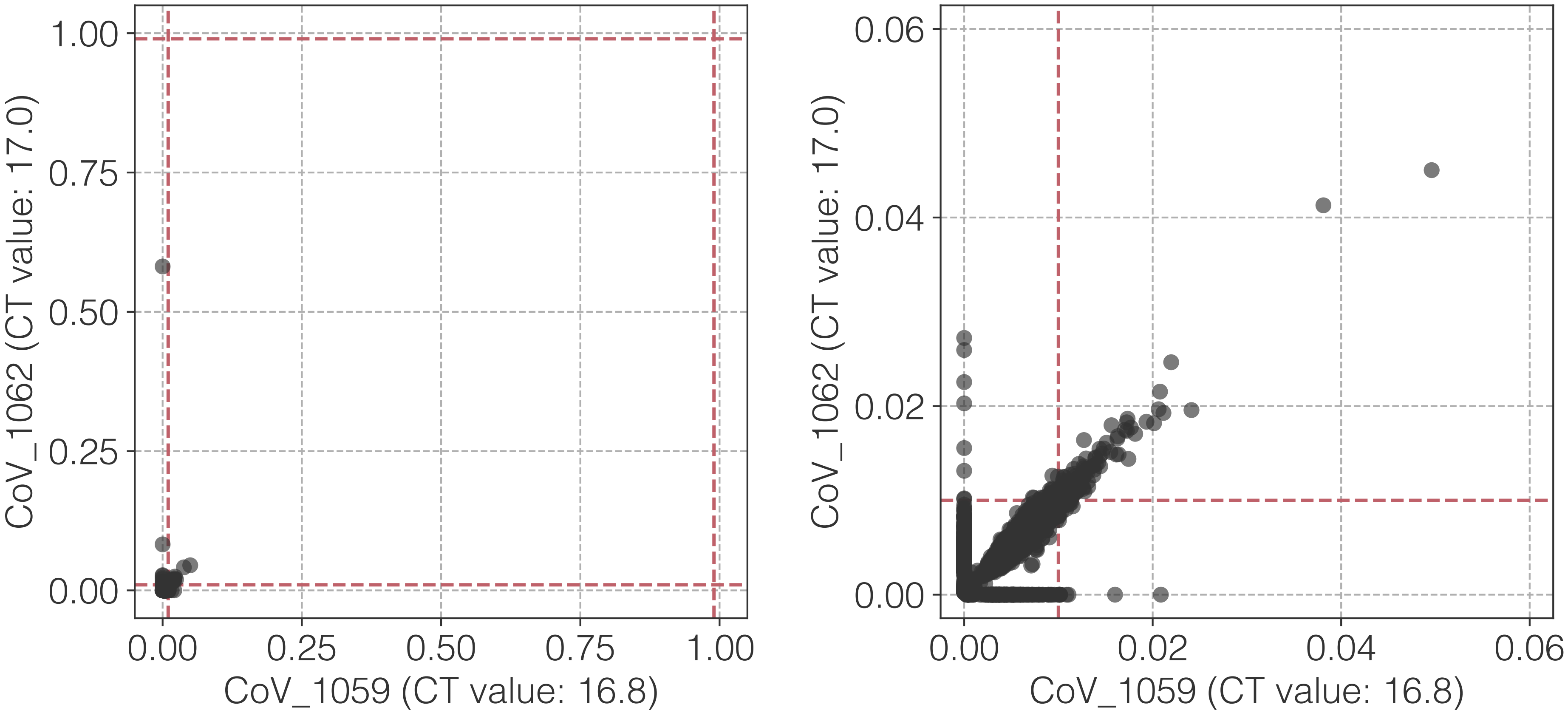

1.00
0.75
0.50
0.25
0.00
CoV_1062 (CT value: 17.0)
0.00
0.25
0.50
0.75
1.00
CoV_1059 (CT value: 16.8)
0.06
0.04
0.02
0.00
CoV_1062 (CT value: 17.0)
0.00
0.02
0.04
0.06
CoV_1059 (CT value: 16.8)

**bb**

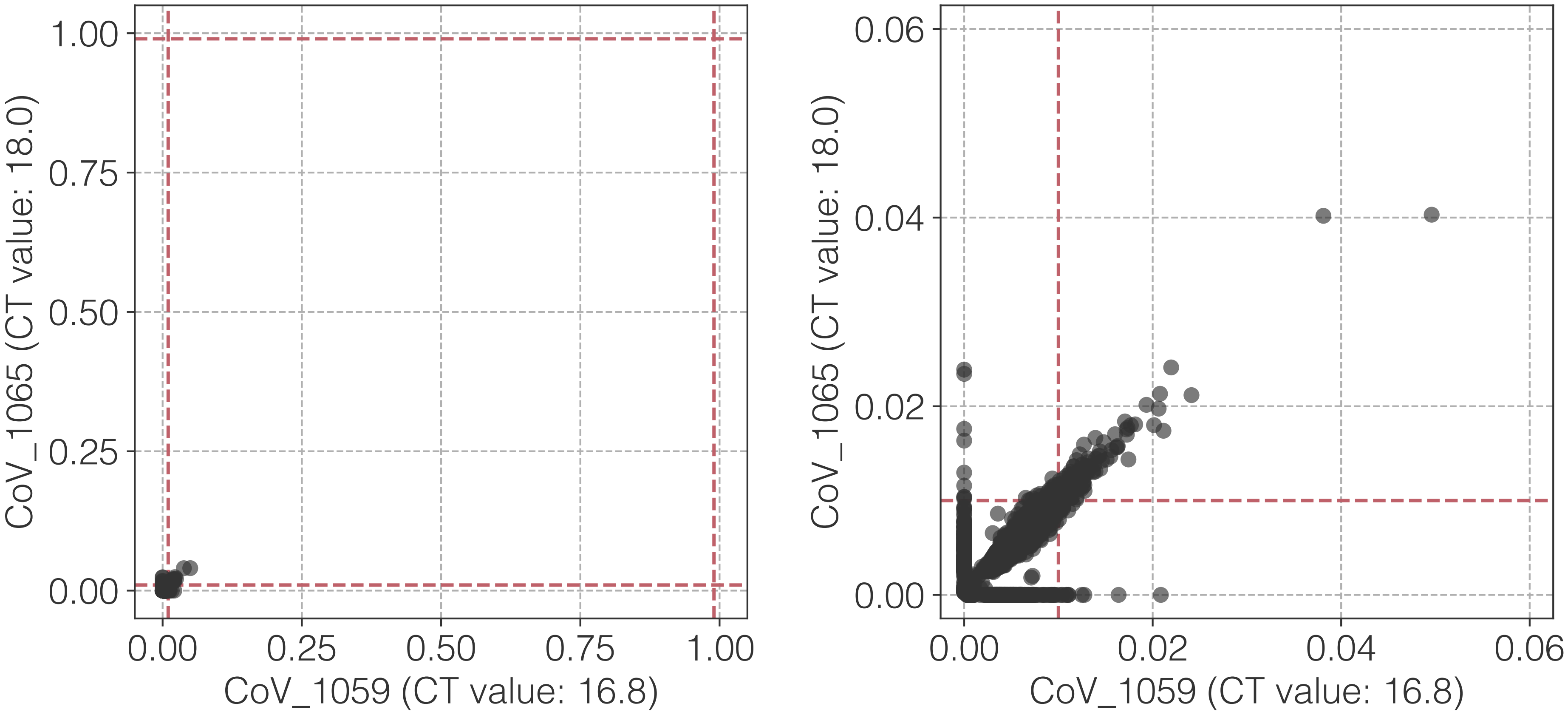

1.00
0.75
0.50
0.25
0.00
0.00
0.25
0.50
0.75
1.00
CoV_1065 (CT value: 18.0)
CoV_1059 (CT value: 16.8)
0.06
0.04
0.02
0.00
0.00
0.02
0.04
0.06
CoV_1065 (CT value: 18.0)
CoV_1059 (CT value: 16.8)

bc

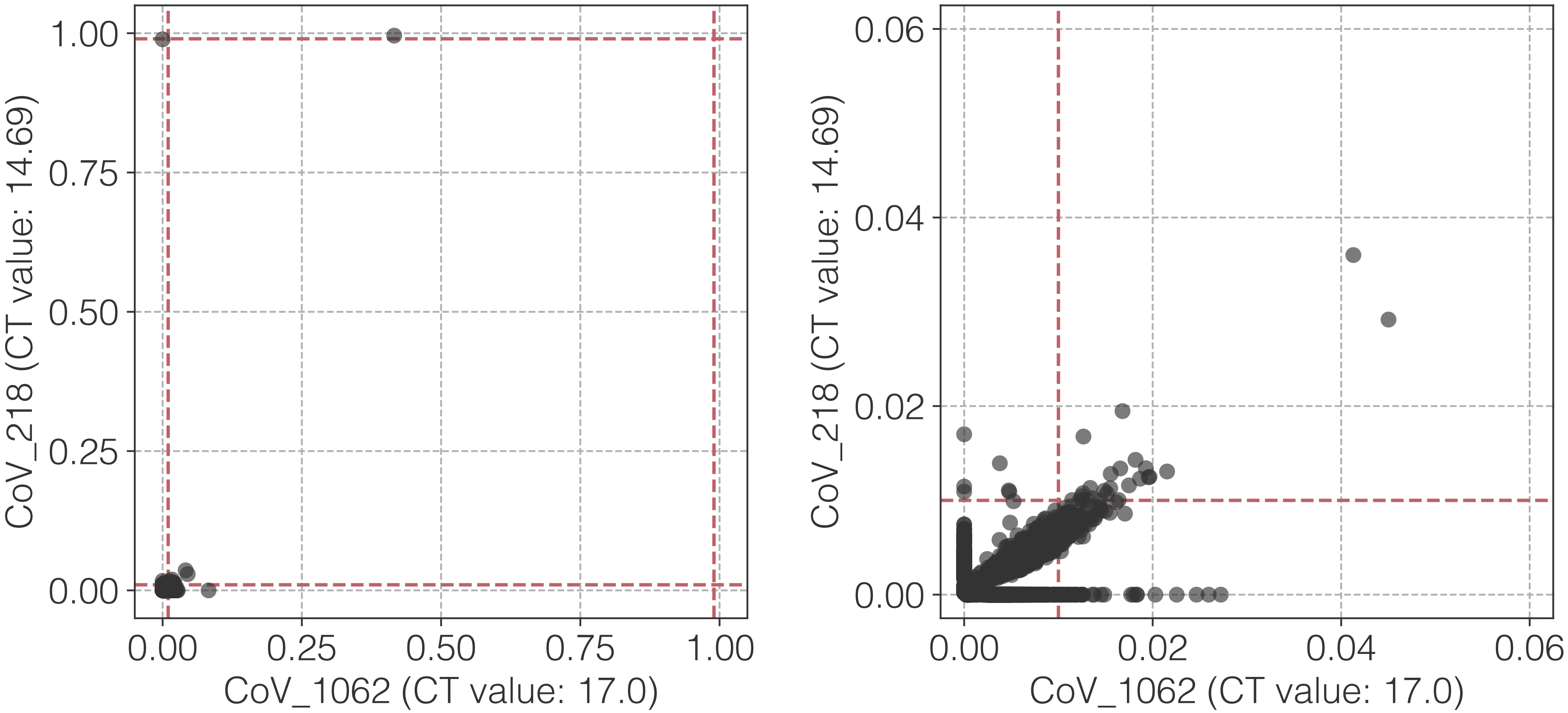

1.00
0.75
0.50
0.25
0.00
CoV_218 (CT value: 14.69)
0.00
0.25
0.50
0.75
1.00
CoV_1062 (CT value: 17.0)
0.06
0.04
0.02
0.00
CoV_218 (CT value: 14.69)
0.00
0.02
0.04
0.06
CoV_1062 (CT value: 17.0)

**bd**

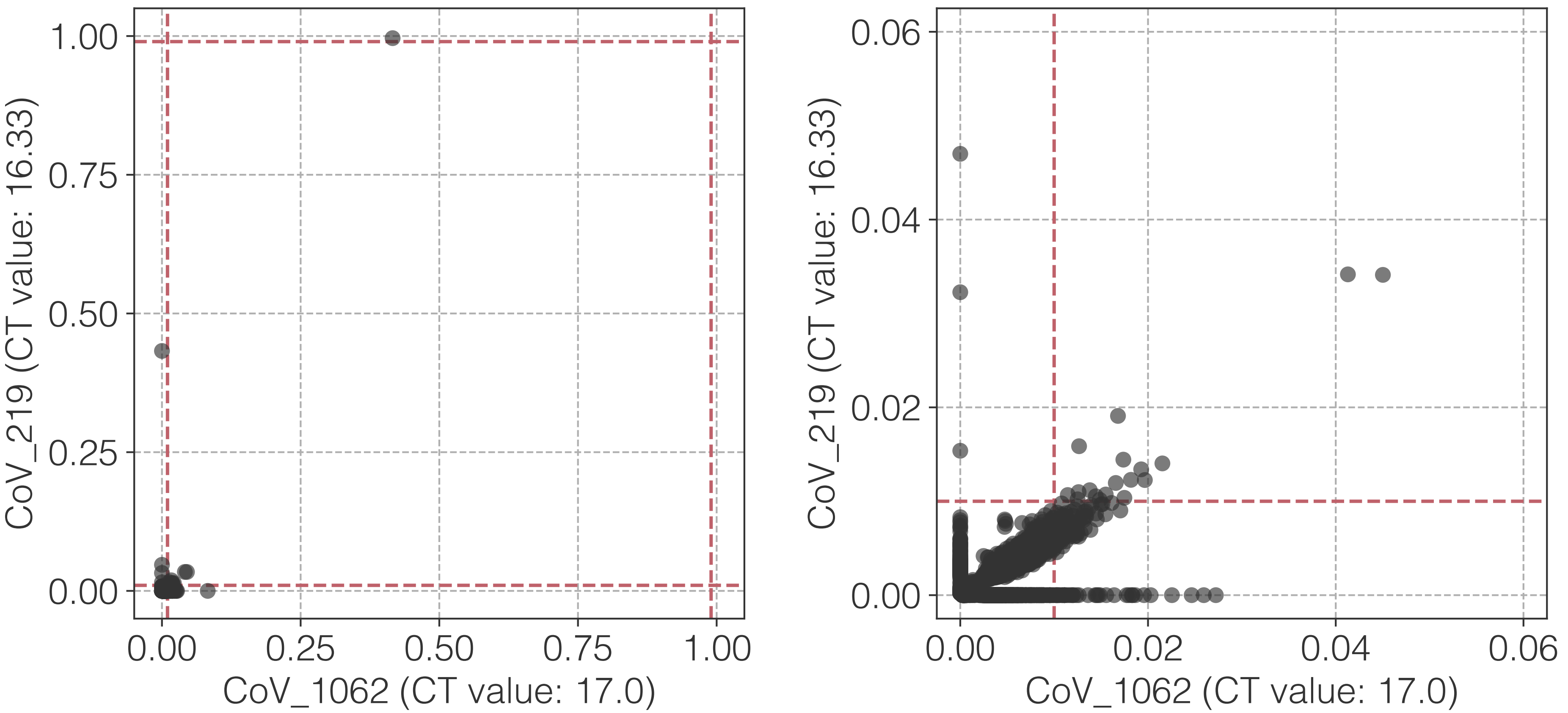
1.00
0.75
0.50
0.25
0.00
CoV_219 (CT value: 16.33)
0.00
0.25
0.50
0.75
1.00
CoV_1062 (CT value: 17.0)
0.06
0.04
0.02
0.00
CoV_219 (CT value: 16.33)
0.00
0.02
0.04
0.06
CoV_1062 (CT value: 17.0)

**be**

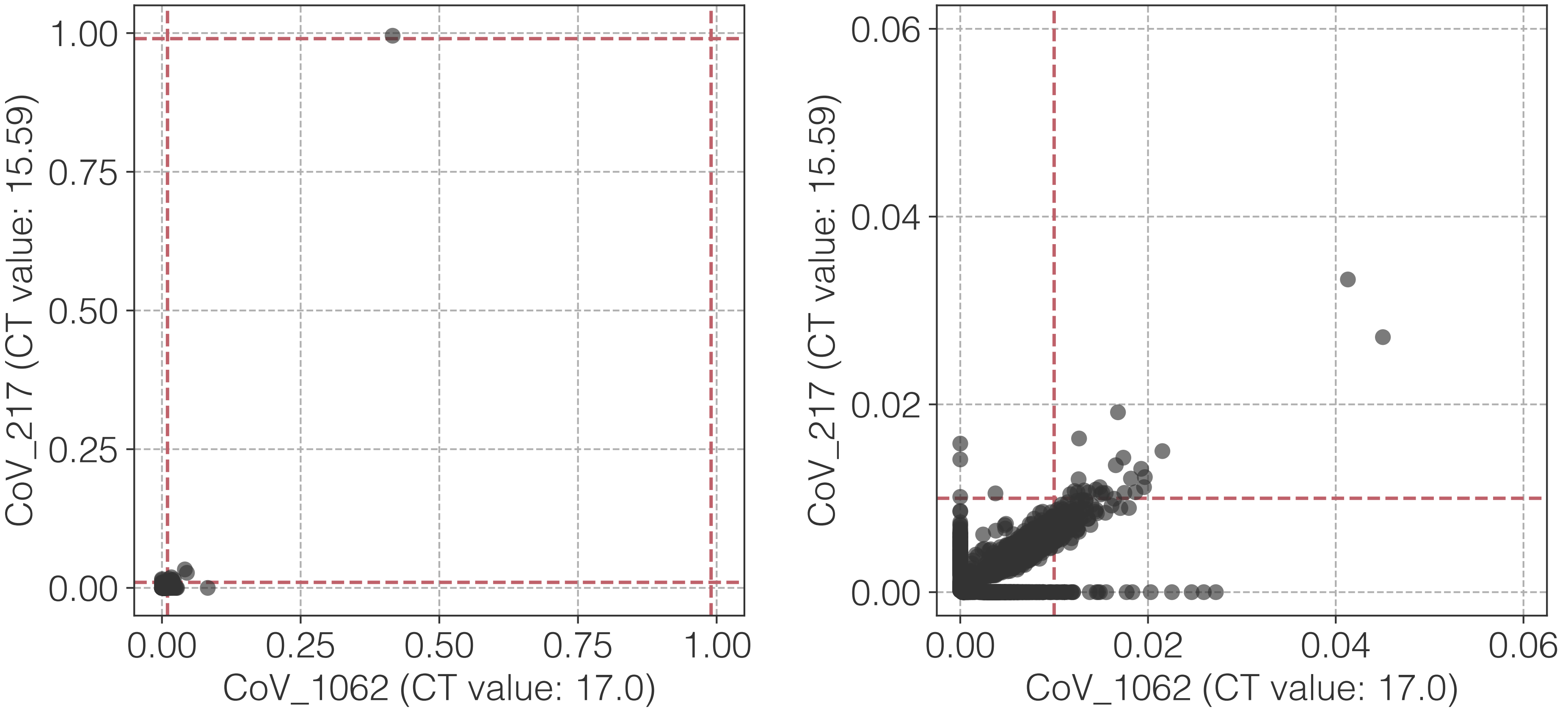

1.00
0.75
0.50
0.25
0.00
CoV_217 (CT value: 15.59)
0.00
0.25
0.50
0.75
1.00
CoV_1062 (CT value: 17.0)
0.06
0.04
0.02
0.00
CoV_217 (CT value: 15.59)
0.00
0.02
0.04
0.06
CoV_1062 (CT value: 17.0)

**bf**

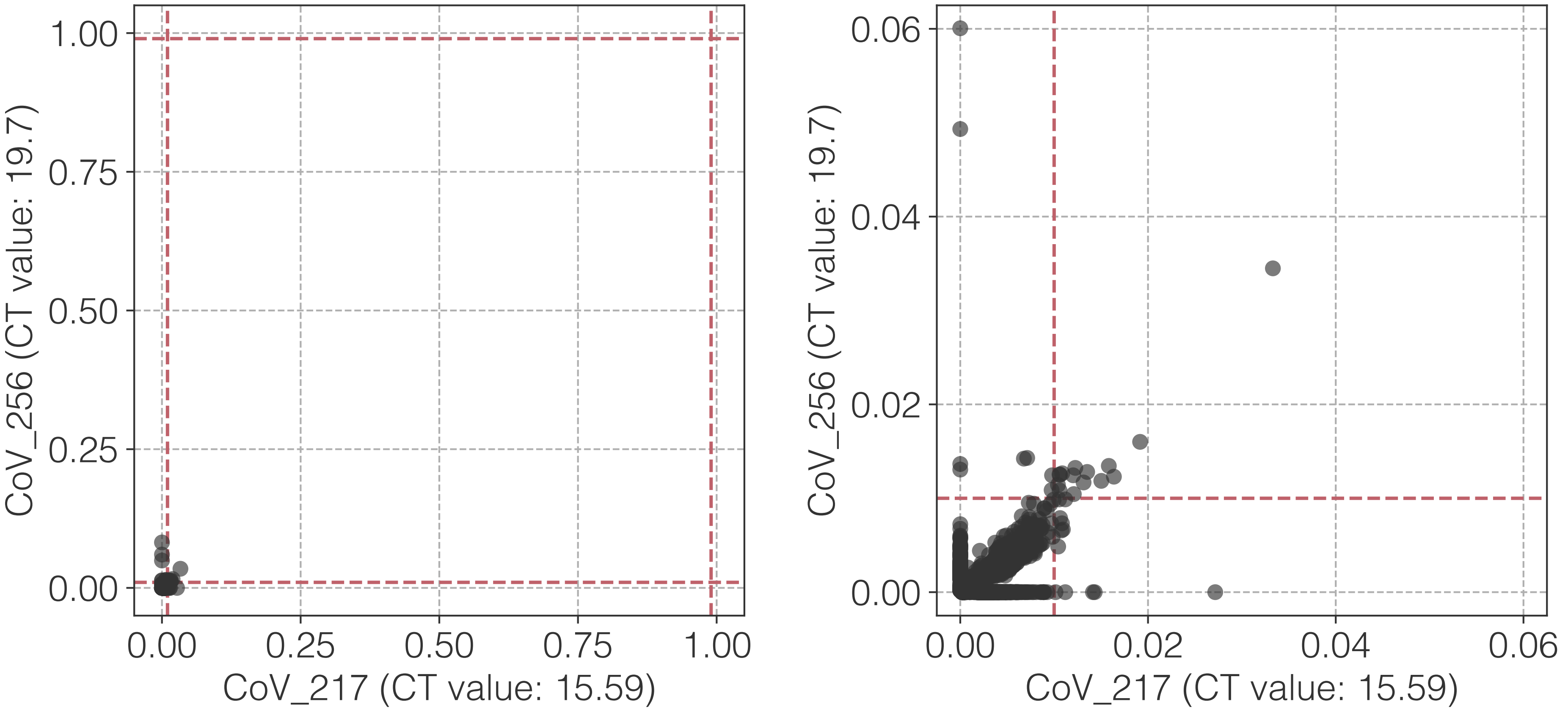

1.00
0.75
0.50
0.25
0.00
CoV_256 (CT value: 19.7)
0.00
0.25
0.50
0.75
1.00
CoV_217 (CT value: 15.59)
0.06
0.04
0.02
0.00
CoV_256 (CT value: 19.7)
0.00
0.02
0.04
0.06
CoV_217 (CT value: 15.59)

**bg**

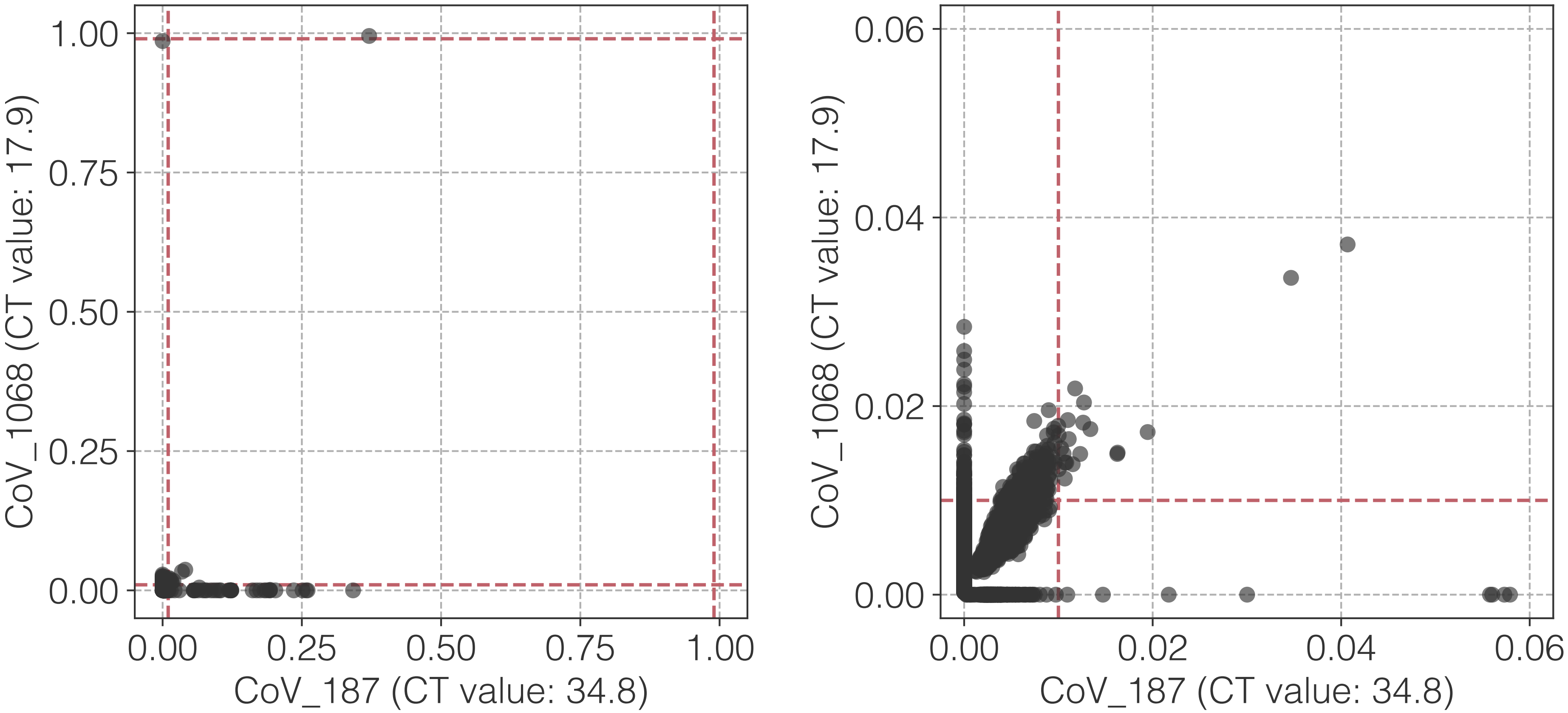

1.00
0.75
0.50
0.25
0.00
CoV_1068 (CT value: 17.9)
0.00
0.25
0.50
0.75
1.00
CoV_187 (CT value: 34.8)
0.06
0.04
0.02
0.00
CoV_1068 (CT value: 17.9)
0.00
0.02
0.04
0.06
CoV_187 (CT value: 34.8)

**bh**

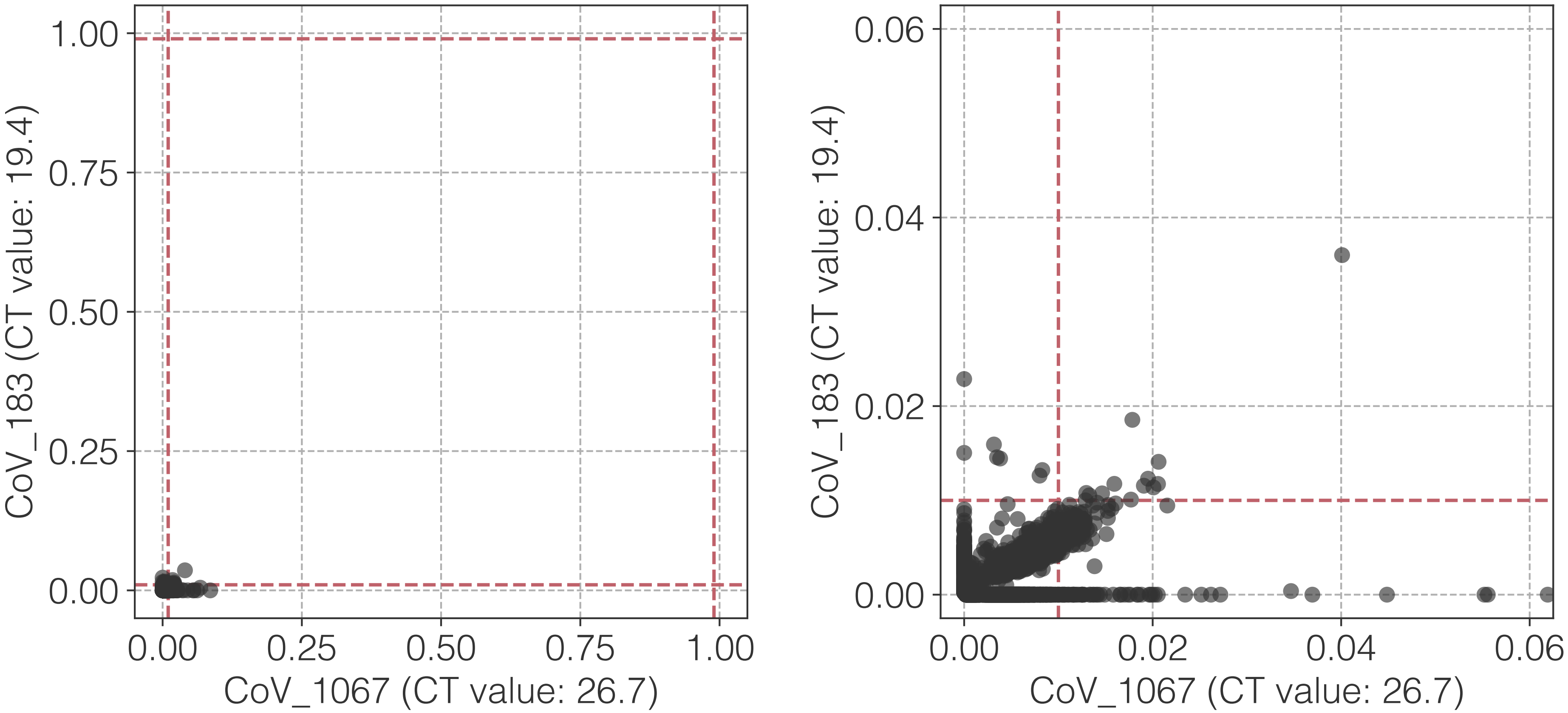
1.00
0.75
0.50
0.25
0.00
CoV_183 (CT value: 19.4)
0.00
0.25
0.50
0.75
1.00
CoV_1067 (CT value: 26.7)
0.06
0.04
0.02
0.00
CoV_183 (CT value: 19.4)
0.00
0.02
0.04
0.06
CoV_1067 (CT value: 26.7)

**bi**

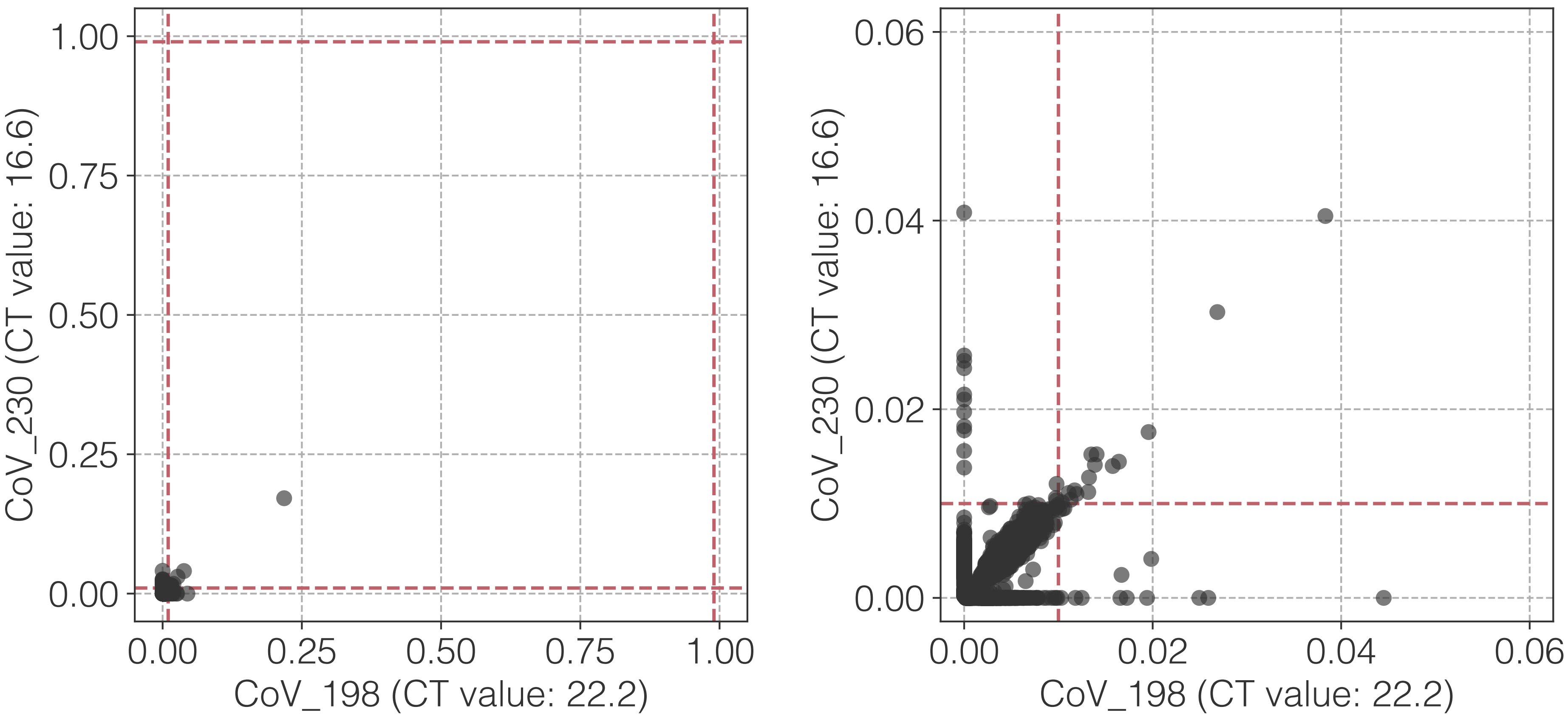

1.00
0.75
0.50
0.25
0.00
CoV_230 (CT value: 16.6)
0.00
0.25
0.50
0.75
1.00
CoV_198 (CT value: 22.2)
0.06
0.04
0.02
0.00
CoV_230 (CT value: 16.6)
0.00
0.02
0.04
0.06
CoV_198 (CT value: 22.2)

**bj**

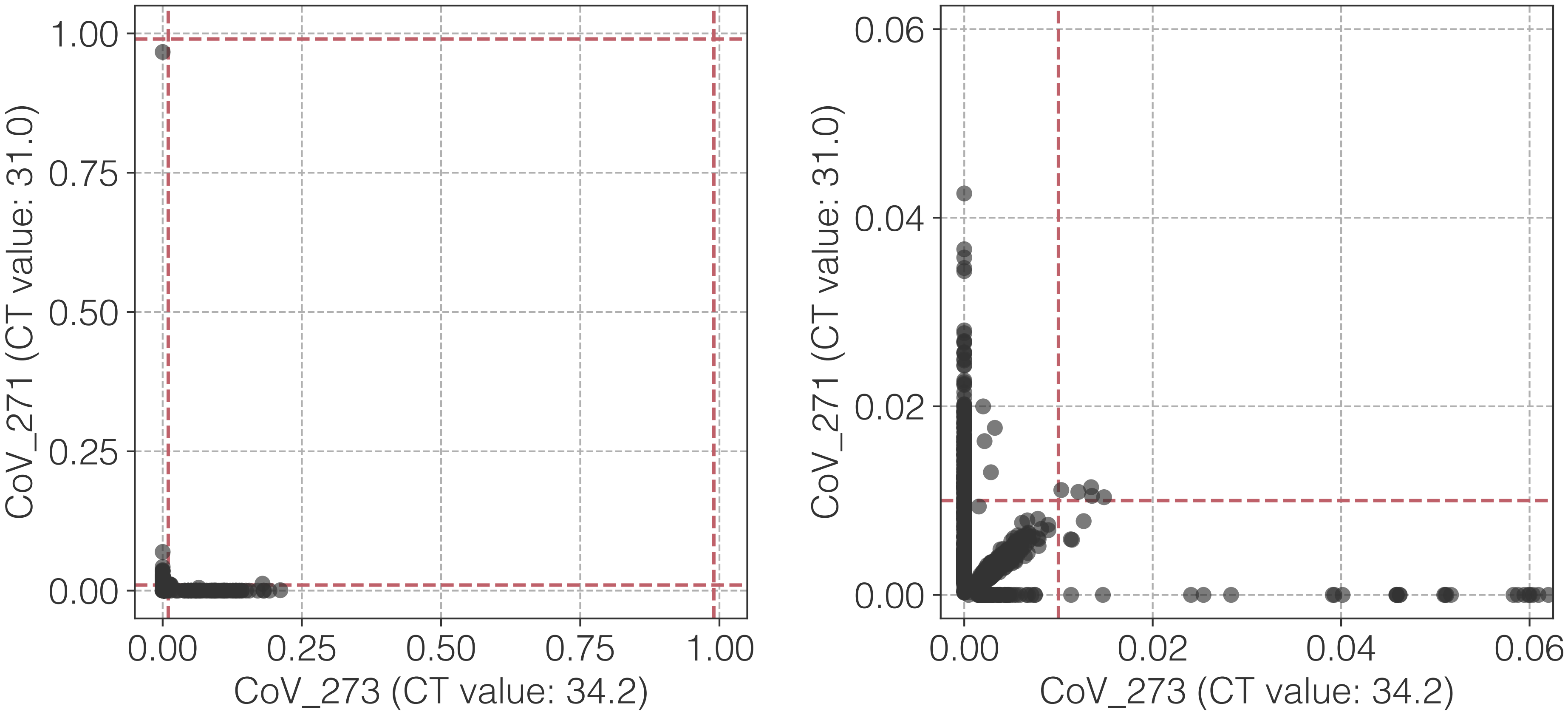

CoV_271 (CT value: 31.0)
CoV_273 (CT value: 34.2)
1.00
0.75
0.50
0.25
0.00
0.06
0.04
0.02
0.00

**bk**

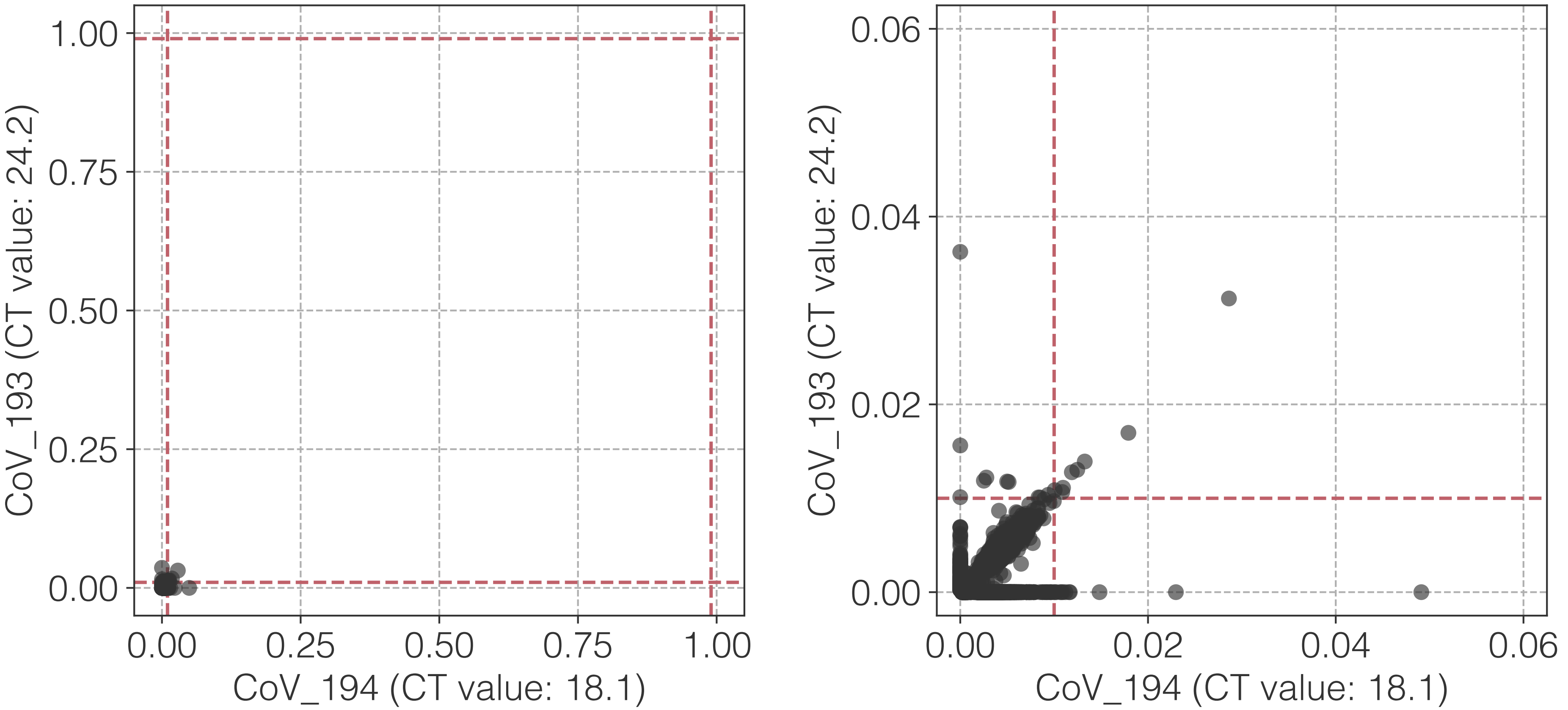

1.00
0.75
0.50
0.25
0.00
CoV_193 (CT value: 24.2)
0.00
0.25
0.50
0.75
1.00
CoV_194 (CT value: 18.1)
0.06
0.04
0.02
0.00
CoV_193 (CT value: 24.2)
0.00
0.02
0.04
0.06
CoV_194 (CT value: 18.1)

**bl**

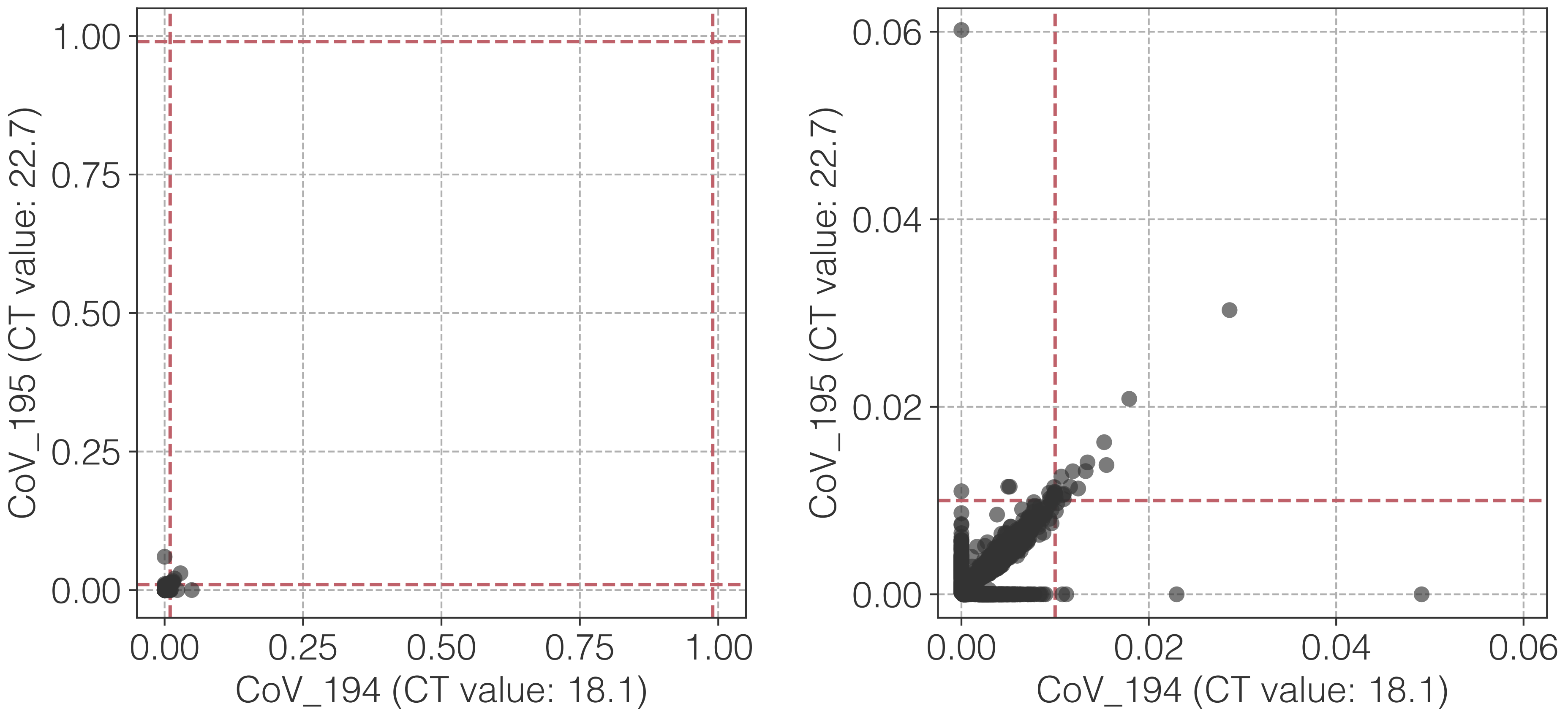

1.00
0.75
0.50
0.25
0.00
CoV_195 (CT value: 22.7)
0.00
0.25
0.50
0.75
1.00
CoV_194 (CT value: 18.1)
0.06
0.04
0.02
0.00
CoV_195 (CT value: 22.7)
0.00
0.02
0.04
0.06
CoV_194 (CT value: 18.1)

**bm**

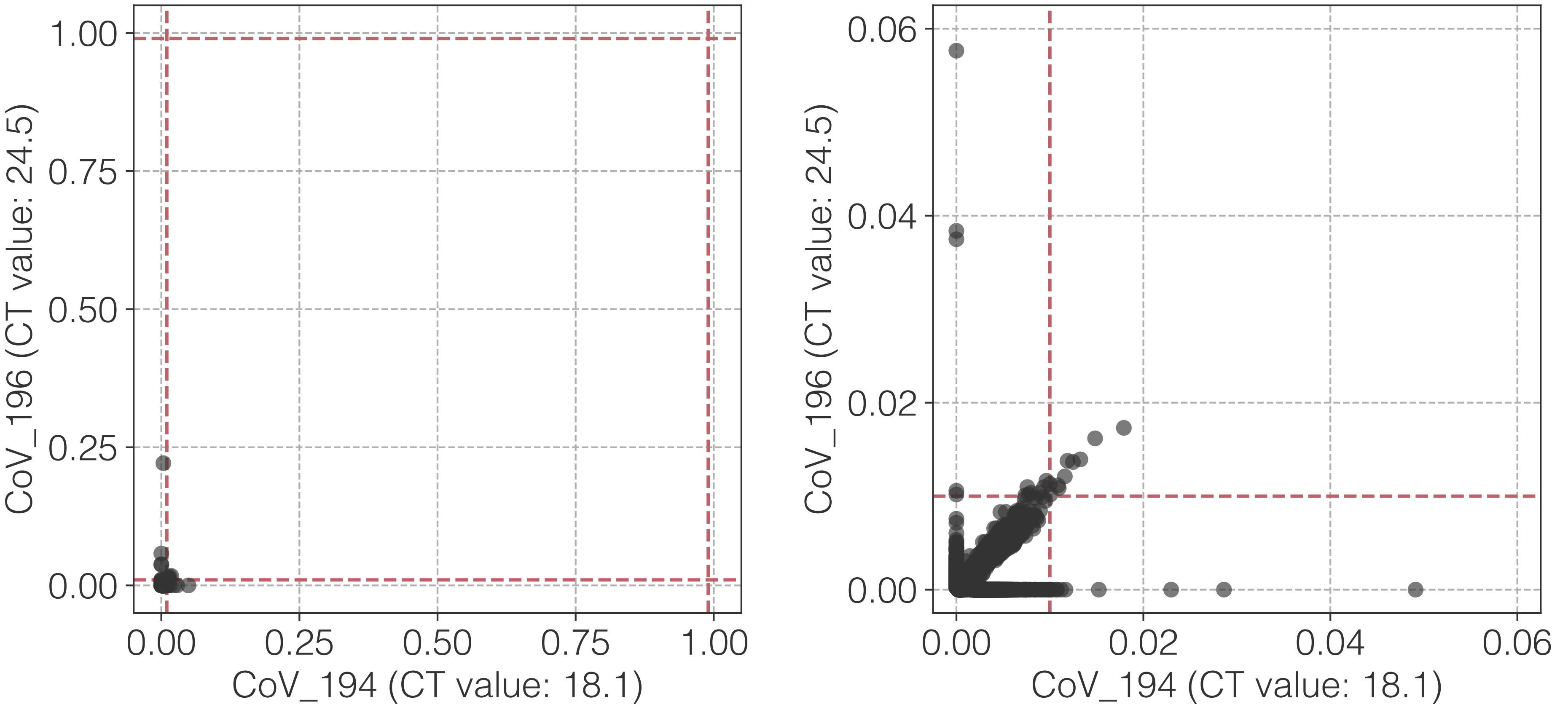
1.00
0.75
0.50
0.25
0.00
CoV_196 (CT value: 24.5)
0.00
0.25
0.50
0.75
1.00
CoV_194 (CT value: 18.1)
0.06
0.04
0.02
0.00
CoV_196 (CT value: 24.5)
0.00
0.02
0.04
0.06
CoV_194 (CT value: 18.1)

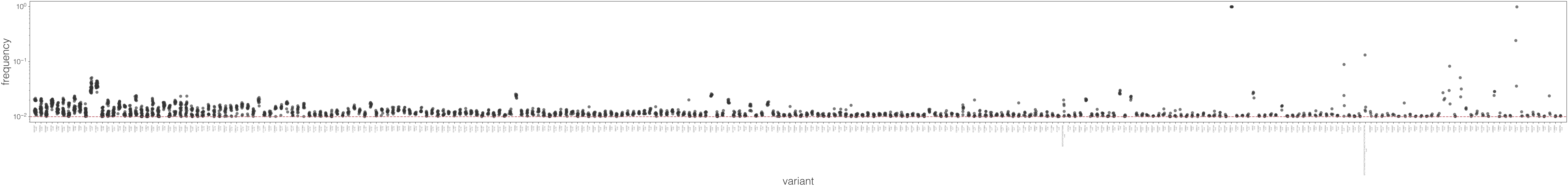

frequency
variant

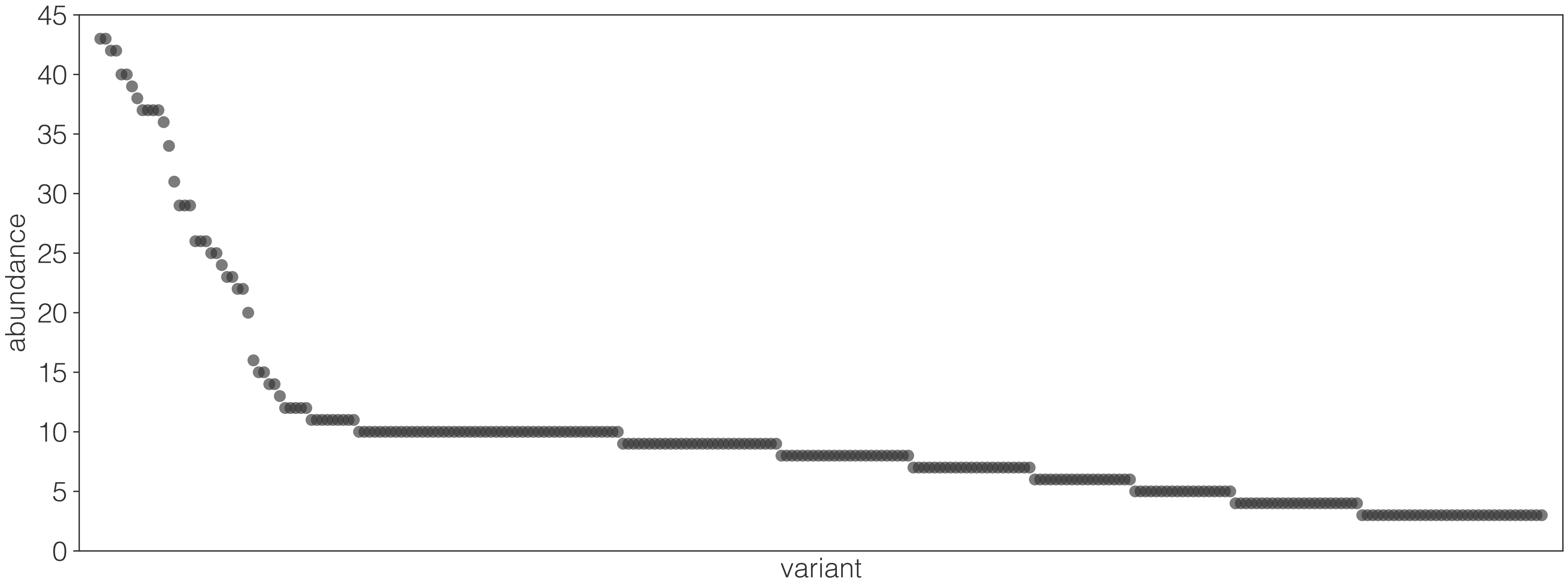

45
40
35
30
25
20
15
10
5
0
abundance
variant
